# Aversive experience drives offline ensemble reactivation to link memories across days

**DOI:** 10.1101/2023.03.13.532469

**Authors:** Yosif Zaki, Zachary T. Pennington, Denisse Morales-Rodriguez, Taylor R. Francisco, Alexa R. LaBanca, Zhe Dong, Sophia Lamsifer, Simón Carrillo Segura, Hung-Tu Chen, Zoé Christenson Wick, Alcino J. Silva, Matthijs van der Meer, Tristan Shuman, André Fenton, Kanaka Rajan, Denise J. Cai

## Abstract

Memories are encoded in neural ensembles during learning and stabilized by post-learning reactivation. Integrating recent experiences into existing memories ensures that memories contain the most recently available information, but how the brain accomplishes this critical process remains unknown. Here we show that in mice, a strong aversive experience drives the offline ensemble reactivation of not only the recent aversive memory but also a neutral memory formed two days prior, linking the fear from the recent aversive memory to the previous neutral memory. We find that fear specifically links retrospectively, but not prospectively, to neutral memories across days. Consistent with prior studies, we find reactivation of the recent aversive memory ensemble during the offline period following learning. However, a strong aversive experience also increases co-reactivation of the aversive and neutral memory ensembles during the offline period. Finally, the expression of fear in the neutral context is associated with reactivation of the shared ensemble between the aversive and neutral memories. Taken together, these results demonstrate that strong aversive experience can drive retrospective memory-linking through the offline co-reactivation of recent memory ensembles with memory ensembles formed days prior, providing a neural mechanism by which memories can be integrated across days.

## Main Text

Individual memories are initially encoded by ensembles of cells active during a learning event^1–5^ and are stabilized during offline periods following learning through reactivation of those ensembles^6–17^. These reactivations often occur in brief synchronous bursts, which are necessary to drive memory consolidation^18–20^. Most research on episodic memory has focused on how the brain maintains stable representations of discrete memories; however, animals are constantly aggregating new memories and updating past memories as new, relevant information is learned^21^. Moreover, most studies of associative learning have focused on cues that directly precede or occur with an outcome. However, oftentimes in nature, a predictor may not immediately precede an outcome but animals are nonetheless capable of learning to make an inference about the association (e.g., conditioned taste aversion)^22^. It is unclear the environmental variables that could promote memories to be linked across long periods (i.e., days), and the neural mechanisms of memory integration across such disparate time periods are poorly understood. In addition, while it has been shown that offline periods support memory consolidation, recent studies have suggested that offline periods following learning may be important for memory integration processes as well^23–26^.

### Strong aversive experience drives retrospective memory-linking

To investigate how memories are integrated across days, we first designed a behavioral experiment to test whether mice would spread fear from an aversive memory to a neutral memory formed two days prior (Retrospective memory-linking) or two days after (Prospective memory-linking) (Figure 1A). In the Retrospective group, mice first experienced a Neutral context followed by an Aversive context paired with a foot shock two days later. In the Prospective group, mice experienced an Aversive context followed by a Neutral context two days later. Both groups were then tested in the Aversive context to test for recall of the aversive memory, followed by testing in the previously experienced Neutral context or an unfamiliar Novel context to test for non-specific fear generalization. Memory-linking was defined as a selective increase in fear in the Neutral context compared to the Novel context, both contexts in which they had never been shocked. Notably, this definition distinguishes memory-linking from a broader generalization of fear across contexts. Mice froze no differently in the Aversive context in either group, suggesting that the perceived negative valence of the Aversive context was not different between groups (Figure 1B). Interestingly, in the Retrospective group, mice froze more in the Neutral context compared to the Novel context, suggesting that fear spread retrospectively from the Aversive context to the Neutral context experienced two days prior. However, in the Prospective group, there was no difference in freezing between the Neutral and Novel contexts, suggesting that memory-linking between the Aversive and Neutral contexts did not occur prospectively across days (Figure 1C). Consistent with prior studies, mice froze in the Neutral context in both Prospective and Retrospective conditions when the Neutral and Aversive contexts were experienced within a day (5h apart, Extended Figure 1A)^27,28^. However, when the contexts were separated by more than one day, mice froze in the Neutral context only in the Retrospective and not the Prospective condition (Extended Figure 1B).

**Figure 1.**
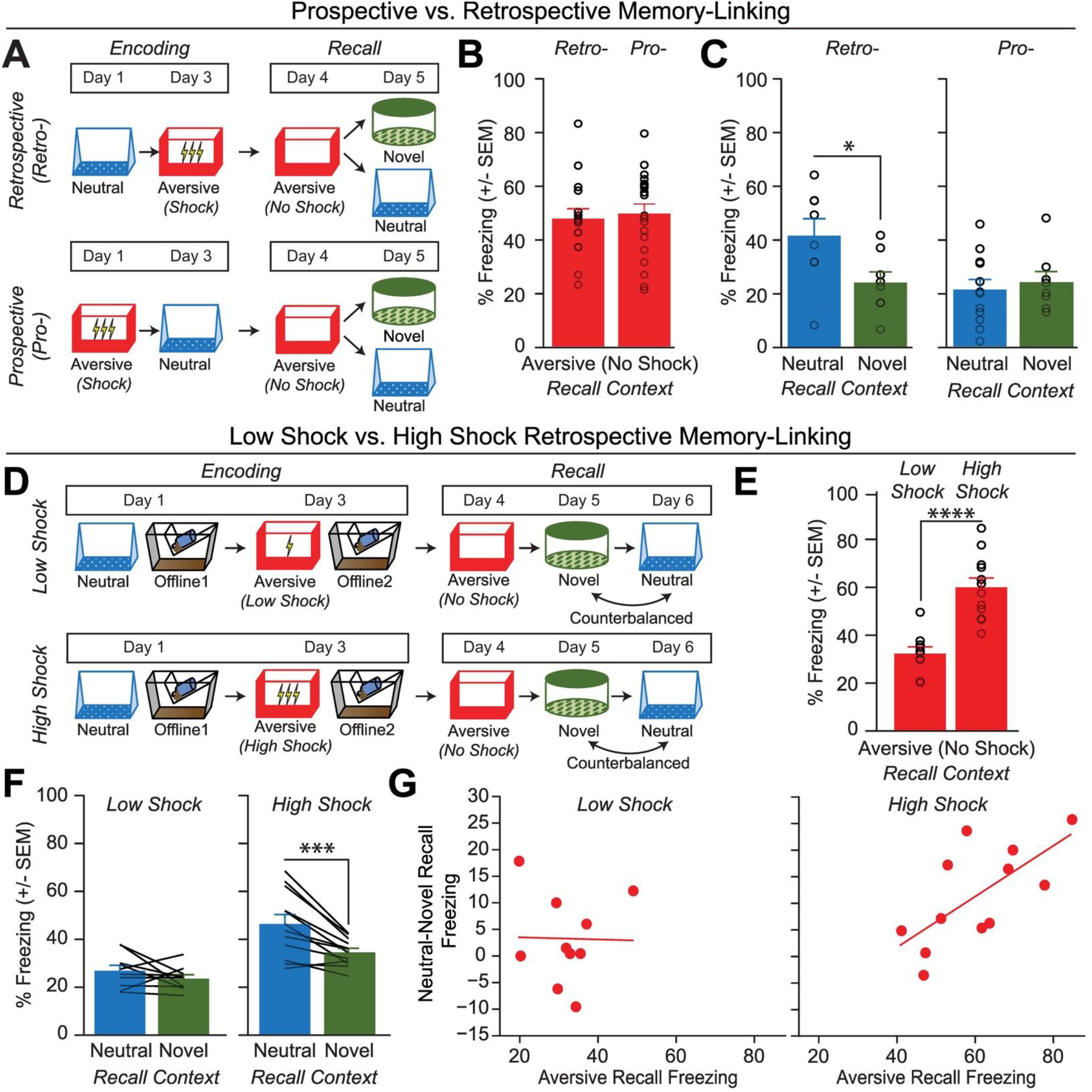
Strong aversive experience drives retrospective memory-linking to a neutral context learned days ago. A) Schematic of prospective vs retrospective memory-linking behavior experiment. Mice either received a Neutral experience followed by an Aversive experience two days later (Retrospective) or the Aversive experience followed by Neutral (Prospective). One day after the second experience, mice were tested in the Aversive context they were shocked in. The following day, mice were tested in either the previously experienced Neutral context or a Novel context. B) Freezing during Aversive recall in Prospective vs Retrospective groups. There was no difference in Aversive recall freezing between Prospective & Retrospective conditions (*t_34_ = 0.36, p = 0.72*) (*Retrospective, N = 16 mice; Prospective, N = 20 mice*). C) Freezing during Neutral vs Novel recall in Prospective vs Retrospective groups. There was a significant interaction between freezing in Neutral vs Novel recall in the Retrospective vs Prospective groups, suggesting the Aversive experience retrospectively linked to the Neutral memory, but not prospectively. Significant interaction between Direction (Prospective vs Retrospective) and Context (Neutral vs Novel), (*F_1,32_ = 4.90, p = 0.034*) (*Retrospective Neutral, N = 8 mice; Retrospective Novel, N = 8 mice; Prospective Neutral, N = 12 mice, Prospective Novel, N = 8 mice*). Post-hoc, *Retrospective (t_32_ = 2.586, p = 0.029*), *Prospective (t_32_ = 0.452, p = 0.6546)*. D) Schematic of Low Shock vs High Shock retrospective memory-linking experiment. Mice received a Neutral experience followed by a 1hr offline session in their homecage. Two days later, they received either 3 low shocks (0.25mA) or 3 high shocks (1.5mA, same amplitude as in Figure 1A) in an Aversive context, followed by another 1hr offline session in their homecage. The following day they were tested in the Aversive context, and for the following two days they were tested in the Neutral and Novel contexts, counterbalanced. Calcium imaging was performed during all the sessions. E) Freezing during Aversive recall in Low vs High Shock mice. Mice froze more in the Aversive context after receiving a high shock vs low shock (*t_18.8_ = 5.877, p = 0.000012*) (*Low Shock, N = 10 mice; High Shock, N = 12 mice*). F) Freezing during Neutral vs Novel recall in Low vs High Shock mice. Mice only displayed enhanced freezing in Neutral vs Novel (i.e., retrospective memory-linking) after High Shock and not Low Shock. Significant effect of Context (Neutral vs Novel) (*F_1,20_ = 17.32, p = 0.000048*) and significant interaction between Context and Amplitude (*F_1,20_ = 4.99, p = 0.037*) (*Low Shock, N = 10 mice; High Shock, N = 12 mice*). High Shock mice froze more in the Neutral vs Novel contexts (*t_11_ = 4.37, p = 0.002*) while Low Shock mice froze no differently in the two contexts (*t_9_ = 1.23, p = 0.249*). G) Correlation between Aversive recall freezing and memory-linking strength. The strength of the aversive memory was correlated with the degree of retrospective memory-linking in High Shock mice (*R_2_ = 0.45, p = 0.016*), but not in Low Shock mice (*R_2_ = 0.0003, p = 0.963*) (*Low Shock, N = 10 mice; High Shock, N = 12 mice*).

We next asked what environmental variables drove two memories to be linked retrospectively across days. It has previously been suggested that the emotional salience of an experience enhances its storage into memory^29,30^, as well as its likelihood of altering past neutral memories in humans^31^. Thus, we hypothesized that the more aversive the experience, the more likely that fear would be retrospectively linked to a previous neutral memory. To test this, we manipulated the shock intensity during aversive encoding to test if stronger shock would drive retrospective memory-linking (Figure 1D). Mice were exposed to a Neutral context followed by an Aversive context paired with a low or high shock two days later (Low Shock group & High Shock group). Mice were then tested in the Aversive, Neutral, and a Novel context in the subsequent three days. As expected, mice in the High Shock group froze more than mice in the Low Shock group during recall in the Aversive context (Figure 1E). We found that only High Shock mice exhibited a selective increase in freezing in the previously experienced Neutral context relative to the Novel context during recall (Figure 1F; Extended Figure 1C-E). If the perceived aversiveness of an experience affects the likelihood of retrospective memory-linking, we hypothesized that levels of freezing during Aversive memory recall would positively correlate with memory-linking— defined as the difference between freezing in the Neutral context and in the Novel context. Indeed, in the High Shock mice, the freezing during Aversive context recall positively correlated with the degree of memory-linking (Figure 1G).

We next investigated how the brain links recent aversive memories with past neutral memories formed days prior. It has been well established in rodents and humans that memories are reactivated during restful periods following learning (i.e., offline periods) to promote the storage of recently learned information^17,32–34^. However, recent work in humans has shown that offline periods can drive the integration of discrete memories as well^23,35,36^. Thus, we hypothesized that following an aversive experience (High Shock group), the offline period may be serving not only to support the consolidation of the aversive memory, but also to link the recent aversive memory with the prior neutral memory, thus increasing freezing during recall of the Neutral context. A major site of memory formation in the brain is the hippocampus, where rapid plasticity following an experience promotes the formation of a memory for that experience and reflects memory expression thereafter^18,27,37–39^. Thus, we asked whether hippocampal activity during the offline period following Aversive encoding was necessary to drive retrospective memory-linking. To do this, we used a chemogenetic manipulation system to disrupt endogenous hippocampal activity during the offline period following Aversive encoding paired with a strong shock (Extended Figure 2). We predicted that this would disrupt retrospective memory-linking. Prior studies have shown that PSAM4-GlyR (PSAM) is an inhibitory ionotropic receptor with no endogenous ligand, and binding of the PSEM ligand with the PSAM receptor causes robust hyperpolarization in neurons^40^. We injected mice with a pan-neuronal, PSAM4-GlyR-expressing virus bilaterally in hippocampus and during the offline period immediately following Aversive encoding, we administered either PSEM to manipulate offline hippocampal activity, or injected saline as a control. We found that mice that received saline during the offline period exhibited a selective increase in freezing in the Neutral over the Novel context, demonstrating retrospective memory-linking. In contrast, mice that received PSEM no longer showed this selective increase in freezing in the Neutral context (Extended Figure 2A-C). To ensure that this effect on retrospective memory-linking was not due to a disrupted memory for the Aversive context, we repeated the experiment, administering PSEM or saline during the offline period, and then tested mice in the Aversive context. We found that mice that received PSEM froze no differently compared to saline controls during Aversive memory recall, suggesting that the strong aversive memory was left intact (Extended Figure 2D,E). These results suggest that hippocampal activity during the offline period is necessary to drive retrospective memory-linking.

### Strong aversive learning drives offline reactivation of a past neutral ensemble

Previous work has suggested that memory reactivation during offline periods following learning could promote not only the consolidation of recently formed memories, but also support the integration of memories^23,25,26,35,36,41^. Consistent with previous studies, we expected that during the offline period following Aversive encoding (while mice are in their homecage), the ensemble active during Aversive encoding would be reactivated to drive consolidation of the recently learned aversive memory. However, we also hypothesized that if the aversive experience was strong enough, the ensemble active during the neutral experience (from two days prior) would be reactivated as well, integrating the neutral and aversive memories.

We first validated that we could detect ensemble reactivation after a salient experience using calcium imaging. To do this, we conducted a contextual fear conditioning experiment, recording hippocampal CA1 calcium dynamics using the open-source UCLA Miniscopes^27^ (Extended Figure 3A,B). We recorded during Aversive encoding, the first hour offline following Aversive encoding, and during recall of the Aversive context and exposure to a Novel context. Consistent with previous literature, we found that the ensemble of cells active during Aversive encoding was reactivated offline and preferentially reactivated during Aversive memory recall, suggesting a stable neural memory ensemble (Extended Figure 3C-K).

To next investigate whether a strong aversive experience was driving offline reactivation of ensembles representing both the aversive and neutral memories, we performed calcium imaging recordings in CA1 during the offline periods following the initial Neutral experience (Offline1) and subsequent Aversive experience (Offline2) in both Low and High Shock groups (Figure 2; Extended Figure 4; same experiment as in Figure 1D). Consistent with the literature^18,20^ and with our previous experiment (Extended Figure 3), following the initial Neutral encoding, the cells that were active during that experience (Neutral ensemble) were more active compared with cells not active during Neutral encoding (Remaining ensemble) in both Low and High Shock groups (Figure 2B, line graphs). There was no difference in the fraction of cells that made up the Neutral ensemble in the Low vs High Shock groups (Figure 2B, pie charts). To measure ensemble reactivation during the offline period after Aversive encoding, we separated cells that were active during the offline period into four ensembles based on when those cells were previously active: Neutral ensemble represented cells active during the initial Neutral encoding and not Aversive encoding; Aversive ensemble represented cells active during Aversive encoding and not Neutral encoding; Neutral ∩ Aversive ensemble represented cells that were active during both Neutral and Aversive encoding; and Remaining ensemble represented cells not observed to be active prior to the offline period (Figure 2C). There was no difference in the fraction of cells that made up each ensemble across Low and High Shock groups (Figure 2C, pie charts). In the Low Shock group, consistent with prior literature^14^, we found the Aversive ensemble, the Neutral ensemble, and the Neutral ∩ Aversive ensemble had higher calcium activity than the Remaining ensemble. And the Neutral ensemble was less active than the Aversive and Neutral ∩ Aversive ensembles (Figure 2C, line graphs, left side). These results are consistent with prior studies demonstrating offline reactivation of neuronal ensembles that were recently active during learning^7–9^. In contrast, in the High Shock group, the Neutral ensemble was no differently active than the Aversive and Neutral ∩ Aversive ensembles (Figure 2C, line graphs, right side), indicating that the high shock increased reactivation of the Neutral ensemble.

**Figure 2.**
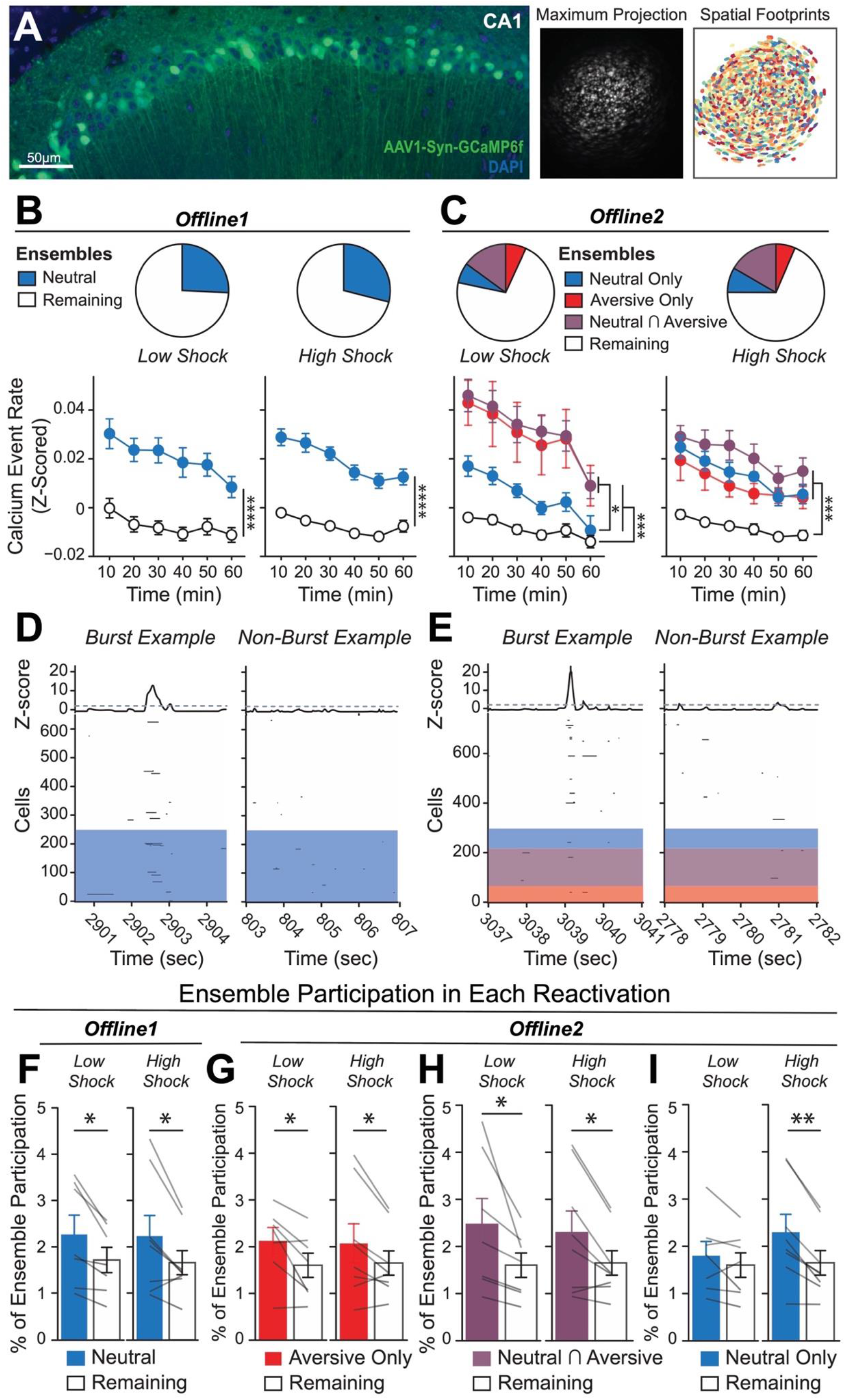
Strong aversive experience drives reactivation of a past neutral ensemble. A) Representative histology (left) of GCaMP6f expression in hippocampal CA1, imaged with a confocal microscope. Green represents AAV1-Syn-GCaMP6f expression, while blue represents a cellular DAPI stain. Maximum intensity projection of an example mouse across one recording session, imaged with a Miniscope (middle), with the spatial footprints of all recorded cells during that session (right) randomly color-coded. B) During Offline1 after Neutral encoding, cells that were active during Neutral encoding (Neutral ensemble) made up ∼25-30% of the offline cell population (pie charts) (*X_2_ = 0.122, df = 1, p = 0.73*). The Neutral ensemble was more highly active than the Remaining ensemble during the offline period (line graphs; A.U.). There was a main effect of Ensemble (*F_1,159_ = 59.19, p = 1.4e-12*), no effect of Amplitude (*F_1,13_ = 0.039, p = 0.85*), and an effect of Time (*F_1,159_ = 4.33, p = 0.039*), and all interactions *p > 0.05* (*Low Shock, N = 7 mice; High Shock, N = 8 mice*). C) During Offline2 after Aversive encoding, similar proportions of previously active cells were reactivated across Low and High shock groups (pie charts) (*X_2_ = 0.326, df = 3, p = 0.955*). However, ensembles were differentially reactivated based upon the amplitude of the Aversive experience (*Ensemble x Amplitude: F_3,331_ = 5.36, p = 0.0013*) (line graphs; A.U.). In Low Shock mice, the Neutral, Aversive, and Neutral ∩ Aversive ensembles were more highly active than the Remaining ensemble (*contrast, t_18_ = 4.22, p = 0.0005*). Additionally, these ensembles were differentially active relative to one another (*F_2,12_ = 4.03, p = 0.046*). This was driven by the Neutral ensemble being less active. The Neutral ensemble was less active than the Aversive and Neutral ∩ Aversive ensembles (*t_12_ = 2.83, p = 0.03*) while the Aversive ensemble was no differently active than the Neutral ∩ Aversive ensemble (*t_12_ = 0.19, p = 0.85*). In High Shock mice, the Neutral, Aversive, and Neutral ∩ Aversive ensembles were all more highly active than the Remaining ensemble (*t_21_ = 4.36, p = 0.0003*), but these three ensembles were no differently active from each other (*F_2,14_ = 1.52, p = 0.25*) (*Low Shock, N = 7 mice; High Shock, N = 8 mice*). D) During the offline periods, hippocampal activity displayed brief bursts of neural activity. To detect these bursts, we computed the z-scored mean activity of the entire recorded population and applied a threshold of z=2 and defined burst periods as all the timepoints above this threshold. The left raster represents an example burst period during Offline1, during which mean population activity briefly reached above threshold. Each row of the raster represents the activity of every recorded neuron, color-coded based on the ensemble it was a part of (blue represents Neutral ensemble and grey represents Remaining ensemble; see legend in Figure 2B). The top black trace represents the z-scored mean population activity. The right raster represents an example non-burst period. E) Same as D but an example burst and non-burst period for Offline2. Each row of the raster again is color-coded based on the ensemble it was a part of (Aversive in red, Neutral ∩ Aversive in purple, Neutral in blue, and Remaining in grey; see legend in Figure 2C). F) During Offline1 in both Low and High Shock groups, a larger fraction of the Neutral ensemble participated in bursts than the Remaining ensemble did (*Ensemble: F_1,13_ = 16.33, p = 0.001; Amplitude: F_1,13_ = 0.009, p = 0.925; Ensemble x Amplitude: F_1,13_ = 0.0058, p = 0.940*) (*Low Shock, N = 7 mice; High Shock, N = 8 mice*). G) During Offline2 in both Low and High Shock groups, a larger fraction of the Aversive ensemble participated in bursts than the Remaining ensemble (*Ensemble: F_1,13_ = 13.57, p = 0.0028; Amplitude: F_1,13_ = 0.000078, p = 0.99; Ensemble x Amplitude: F_1,13_ = 0.16, p = 0.69*) (*Low Shock, N = 7 mice; High Shock, N = 8 mice*). H) During Offline2 in both Low and High Shock groups, a larger fraction of the Neutral ∩ Aversive ensemble participated in bursts than the Remaining ensemble (*Ensemble: F_1,13_ = 13.95, p = 0.0025; Amplitude: F_1,13_ = 0.014, p = 0.91; Ensemble x Amplitude: F_1,13_ = 0.31, p = 0.58*) (*Low Shock, N = 7 mice; High Shock, N = 8 mice*). I) During Offline2, Neutral and Remaining ensembles differentially participated in bursts in High and Low Shock groups (*Ensemble x Amplitude: F_1,13_ = 5.186, p = 0.040*). High Shock mice showed higher participation of the Neutral ensemble relative to Remaining ensemble (*t_7_ = 4.88, p = 0.0036*), whereas Low Shock mice showed no different participation between the two ensembles (*t_6_ = 1.33, p = 0.23*) (*Low Shock, N = 7 mice; High Shock, N = 8 mice*).

Since the Neutral ensemble was more highly reactivated after high shock, we next investigated whether the Neutral, Aversive, and Neutral ∩ Aversive ensembles might be firing together on a finer temporal scale. Hippocampal activity is known to exhibit organized bursts, oftentimes accompanied by sharp-wave ripples in the local field potential, during which cells active during learning are preferentially reactivated^18^. These events have been found to support memory consolidation^18–20^. Although calcium dynamics are of a coarser timescale than sharp-wave ripples, we observed that during the offline recordings, hippocampal calcium activity periodically exhibited brief bursts of activity during which numerous cells were co-active (Extended Figure 5A,B, from our validation study in Extended Figure 3), consistent with previous reports^42,43^. We found that these bursts were unlikely to occur from shuffled neuronal activities, suggesting that these events were organized events during which groups of hippocampal neurons were synchronously active (Extended Figure 5C-F). We isolated these brief burst periods to ask whether ensembles that were previously active during encoding were selectively participating in these brief burst events (Figure 2D-I; Extended Figure 5A,B; see Methods). We first measured these burst events after a single Aversive learning experience and found that a larger fraction of Aversive ensemble cells participated in these events than the Remaining ensemble cells (Extended Figure 5L). Interestingly, these burst events coincided with the mouse briefly slowing down about 1 second prior to the event, and about 1 second after its onset resuming its locomotion, suggesting that these bursts occurred during periods of brief quiescence (Extended Figure 5I,J)^18^.

We then asked whether a strong shock paired with an Aversive context would drive the Neutral ensemble to also participate within these bursts after Aversive encoding (experiment from Figure 1D). In both Low and High Shock mice and after both Neutral and Aversive encoding, frequencies of burst events (defined by periods when the mean activity of the entire recorded population reached above a required threshold; see Methods) were comparable across groups and decreased across the hour (Extended Figure 4G,H). As expected, after Neutral encoding, both Low and High Shock groups had a larger fraction of the Neutral ensemble participating in these burst events than the Remaining ensemble (Figure 2D,F). After Aversive encoding, both groups again showed selective participation of the Aversive ensemble that was most recently active (Figure 2G) as well as of the Neutral ∩ Aversive ensemble that was previously active during both learning events (Figure 2H). However, only in the High Shock group (and not the Low Shock group) the Neutral ensemble selectively participated in these burst events as well (Figure 2I), suggesting that a strong aversive experience drove the recruitment of the Neutral ensemble into these burst events.

### Strong aversive experience drives co-bursting of the Neutral ∩ Aversive ensemble with the Neutral ensemble

Since after High Shock, the Neutral and Aversive ensembles were both participating in burst events, we next asked whether the two ensembles co-participated within the same bursts, or whether they participated separately in different bursts. Co-bursting between the Neutral ensemble and Aversive ensemble could suggest a process through which the two ensembles can become integrated into a cell assembly likely to co-fire during memory recall thereafter. This process could occur through Hebbian plasticity^44^ or through behavioral timescale synaptic plasticity, which has been proposed to drive the formation of place fields in hippocampal neurons^37^. Previous work has shown that hippocampal neurons become highly co-active during recall of an aversive memory but not during initial learning^45^, that co-activity relationships among hippocampal neurons can distinguish between contexts that a mouse has experienced^46^, and that ensembles that are highly co-active during an offline period following learning are more likely to be reactivated during memory recall than non-co-active neurons^15^.

To ask whether the Neutral, Aversive, and Neutral ∩ Aversive ensembles were co-bursting after Aversive encoding, we measured the fraction of burst events that each ensemble participated in independently of each other (Figure 3A) and the fraction that the ensembles co-participated in (Figure 3D) during the offline period following the Aversive experience (Extended Figure 4I; see Methods).

Previously, we had found that the Neutral ∩ Aversive cells (those active during both Neutral and Aversive encoding) were the most highly active during the offline period (Figure 2C). Highly active subpopulations of neurons have been proposed to form a ‘hub-like’ population of neurons that may orchestrate the activity of other neurons in a larger network^47,48^. Therefore, these highly active neurons could be organizing the activity of other neurons in the hippocampus to drive activity during this offline period. Thus, we hypothesized that co-participation between the highly active Neutral ∩ Aversive ensemble and the Neutral ensemble would be enhanced after a strong aversive experience.

We found that during burst events, the Neutral ∩ Aversive ensemble participated independently more frequently than the Neutral and Aversive ensembles did, but there was no difference between Low and High Shock mice (Figure 3B). Notably, during non-burst periods, independent ensemble bursting did not vary between any of the ensembles (Figure 3C). We next measured co-participation of the ensembles in all combinations (Figure 3D). We found that in the Low Shock group, co-participation between the three ensembles was less likely to occur than the other combinations; however, surprisingly, in the High Shock group, co-participation between the three ensembles was no different from the other combinations (Figure 3E). Additionally, in the High Shock group, the Neutral ∩ Aversive ensemble co-participated with the Neutral ensemble more than it did with the Aversive ensemble, whereas in the Low Shock group, the Neutral ∩ Aversive ensemble co-participated no differently with the Neutral and Aversive ensembles (Figure 3E). Importantly, there were no differences in ensemble co-bursting between Low and High Shock groups during non-burst periods (Figure 3F), suggesting that the ensemble co-participation was confined to periods when the hippocampus was synchronously active. These results suggested that after a strong aversive experience, the Neutral ∩ Aversive ensemble was preferentially co-bursting with the Neutral ensemble. To confirm that this was the case, we used cross-correlations as another measure of co-activity to measure how co-active the Neutral ∩ Aversive ensemble was with the Neutral and the Aversive ensembles. Indeed, only in the High Shock group, the Neutral ∩ Aversive ensemble was preferentially correlated with the Neutral ensemble compared with the Aversive ensemble during the offline period (Extended Figure 4K). Collectively, these results suggest that a strong aversive experience increases the co-bursting of the Neutral ∩ Aversive ensemble with the Neutral ensemble, perhaps to link fear of the recent aversive experience with the past neutral memory.

### Strong aversive experience drives co-reactivation of the Neutral & Aversive and Neutral ensembles during Neutral context recall

Finally, we asked whether hippocampal ensemble reactivation could support the freezing observed in the Neutral context during recall after a high shock and not low shock (as shown in Figure 1F). To do this, we measured hippocampal ensemble activity while mice recalled the Neutral context after the offline period, compared with ensemble activity when they were placed in a Novel context as a control (Figure 4A). Since High Shock mice froze significantly more in the Neutral vs Novel contexts during recall (Figure 1F), we hypothesized that Neutral context recall would drive the aversive memory representation to be reactivated, whereas exposure to a Novel context would not provoke the reactivation of the aversive memory representation. Previously, we found that during the offline period, the Neutral ∩ Aversive ensemble specifically co-reactivated with the Neutral ensemble (Figure 3E, Extended Figure 4K), perhaps forming an integrated ensemble of neurons that is more likely to fire together in the future. If this were the case, when High Shock mice recalled the Neutral context and reactivated the Neutral ensemble, we predicted they might also reactivate the Neutral ∩ Aversive ensemble, perhaps through a process of pattern completion^49^, thereby driving freezing in the Neutral context. Importantly, we expected this not to occur in Low Shock mice, where Neutral and Neutral ∩ Aversive ensemble co-reactivation was not observed, or in High Shock mice during Novel context exposure, since fear did not selectively spread to the Novel context (Figure 1F).

During recall of the Neutral context and exposure to a Novel context, we measured the fraction of cells active during that session which were previously active during encoding of the Neutral or Aversive contexts or active during both Neutral and Aversive encoding (Figure 4A; Extended Figure 4E,F). We previously observed that during the offline period, the Neutral ensemble co-fired with the Neutral ∩ Aversive after high shock but not after low shock (Figure 3D,E; Extended Figure 3K), potentially forming an integrated ensemble that is more likely to fire together later. Thus, we hypothesized that after high shock, during Neutral context recall, the Neutral ensemble (representing the Neutral context) would be reactivated, and this would, in turn, trigger reactivation of the Neutral ∩ Aversive ensemble. As expected, cells exclusively active during Neutral encoding and not Aversive encoding were more likely to be reactivated during Neutral recall than during Novel context exposure in both Low Shock and High Shock groups, suggesting a stable and selective neural population representing the neutral memory (Figure 4B). The cells exclusively active during Aversive encoding were not selectively reactivated during Neutral or Novel contexts in either group (Figure 4C). Interestingly, the cells active during both Neutral and Aversive encoding (Neutral ∩ Aversive ensemble) were more reactivated during Neutral recall than during Novel context exposure in the High Shock but not the Low Shock group (Figure 4D). This suggests that after ensemble co-reactivation during the offline period following high shock, the Neutral ensemble and the Neutral ∩ Aversive ensembles were more likely to reactivate together during Neutral recall.

**Figure 3.**
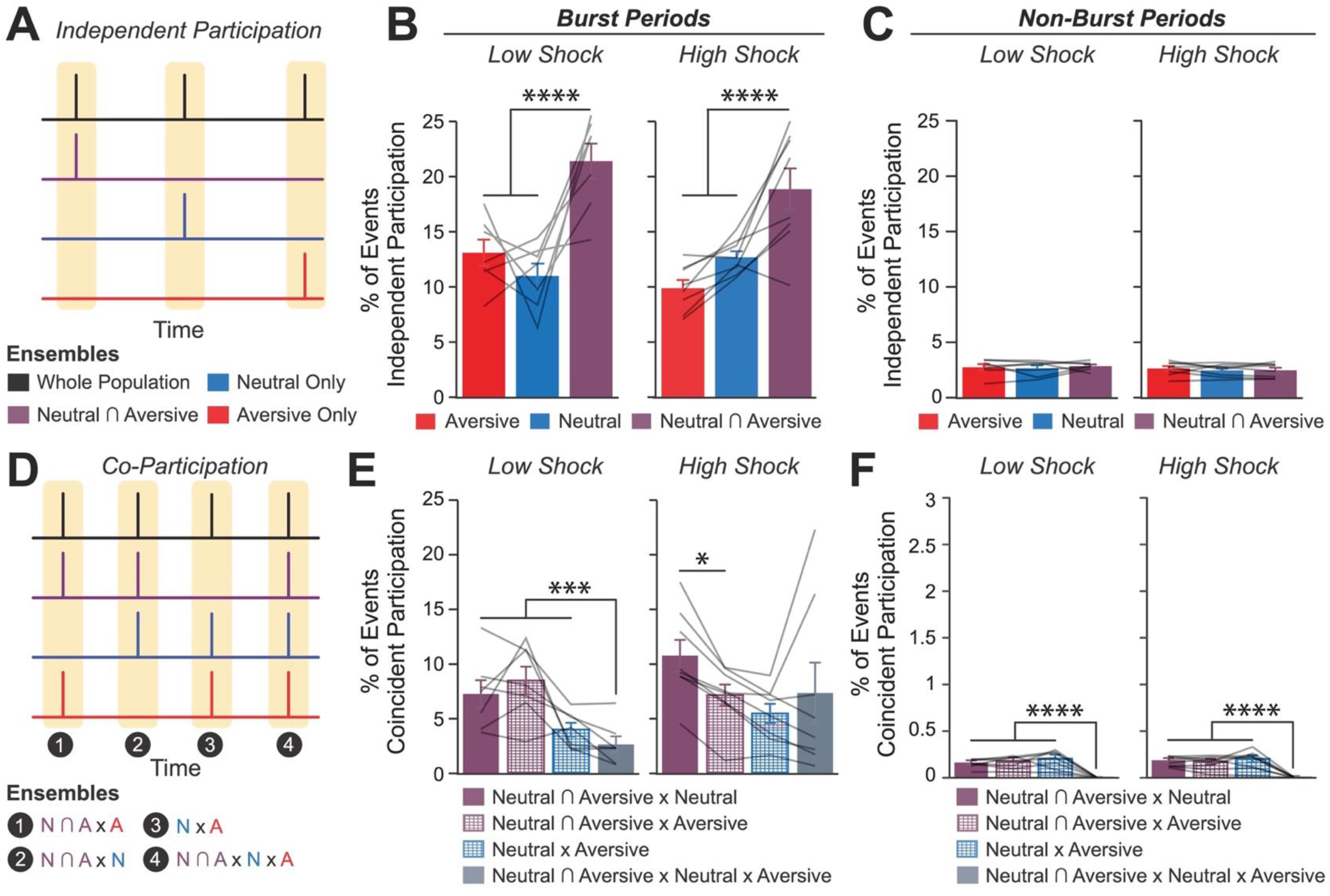
Strong aversive experience drives co-reactivation of the Neutral ensemble with the Neutral ∩ Aversive ensemble. A) Representation of the quantification of independent participation during bursts versus non-bursting periods. Burst events were defined by the whole recorded population, as in Figure 2E (outlined by yellow rectangles). However, now the z-scored mean population activity of the Aversive, Neutral, and Neutral ∩ Aversive ensembles was computed to ask how frequently each ensemble participated in whole population bursts independently of one another. Independent participation meant one ensemble participated while the other two did not. B) During burst periods, the Neutral ∩ Aversive ensemble participated independently in more bursts than the Aversive ensemble (*t_14_ = 7.95, p = 0.000002*) and more than the Neutral ensemble (*t_14_ = 5.59, p = 0.0001*) but there was no difference in participation across Low vs High Shock mice (*F_1,13_ = 1.43, p = 0.25*) and no interaction (*F_2,26_ = 2.49, p = 0.10*) (*Low Shock, N = 7 mice; High Shock, N = 8 mice*). C) During non-burst periods, there was no difference in participation across ensembles (*F_2,26_ = 0.38, p = 0.69*) or between Low and High Shock mice (*F_1,13_ = 0.73, p = 0.41*), and no interaction (*F_2,26_ = 0.36, p = 0.70*) (*Low Shock, N = 7 mice; High Shock, N = 8 mice*). D) Representation of the quantification of co-participation during bursts vs non-bursting periods. As in Figure 3B, the whole population was used to define bursts and the z-scored mean population activities were used to define participation of each ensemble. Co-participation was defined as a whole population burst (outlined by yellow rectangles) during which multiple ensembles participated simultaneously. There were four possible combinations (from left to right: N∩A x N, N∩A x A, N x A, N∩A x N x A) (N∩A = Neutral ∩ Aversive; N = Neutral; A = Aversive). E) During burst periods, there was a significant interaction between Ensemble Combination and Low vs High Shock (*p = 0.01*), suggesting that the patterns of co-bursting varied in Low vs High Shock mice. Post-hoc tests revealed that in Low Shock mice, co-participation between all 3 ensembles was less likely to occur than the other combinations (*t_18_ = 4.73, p = 0.0003*), while in High Shock mice, co-participation between all 3 ensembles occurred no differently than the other combinations (*t_21_ = 0.358, p = 0.72*). Additionally, in the High Shock group, the N∩A ensemble preferentially co-participated with the Neutral ensemble compared to with the Aversive ensemble (*t_21_ = 2.373, p = 0.05*), whereas in the Low Shock group, the N∩A ensemble participated no differently with the Neutral and Aversive ensembles (*t_18_ = 1.196, p = 0.25*) (*Low Shock, N = 7 mice; High Shock, N = 8 mice*). F) During non-burst periods, co-participation between all 3 ensembles was less likely than the other combinations (*t_39_ = 10.92, p = 1.98e-13*); however, there was no effect of Low vs High Shock (*F_1,13_ = 0.038, p = 0.847*) and no interaction (*F_3,39_ = 0.198, p = 0.897*) (*Low Shock, N = 7 mice; High Shock, N = 8 mice*).

**Figure 4.**
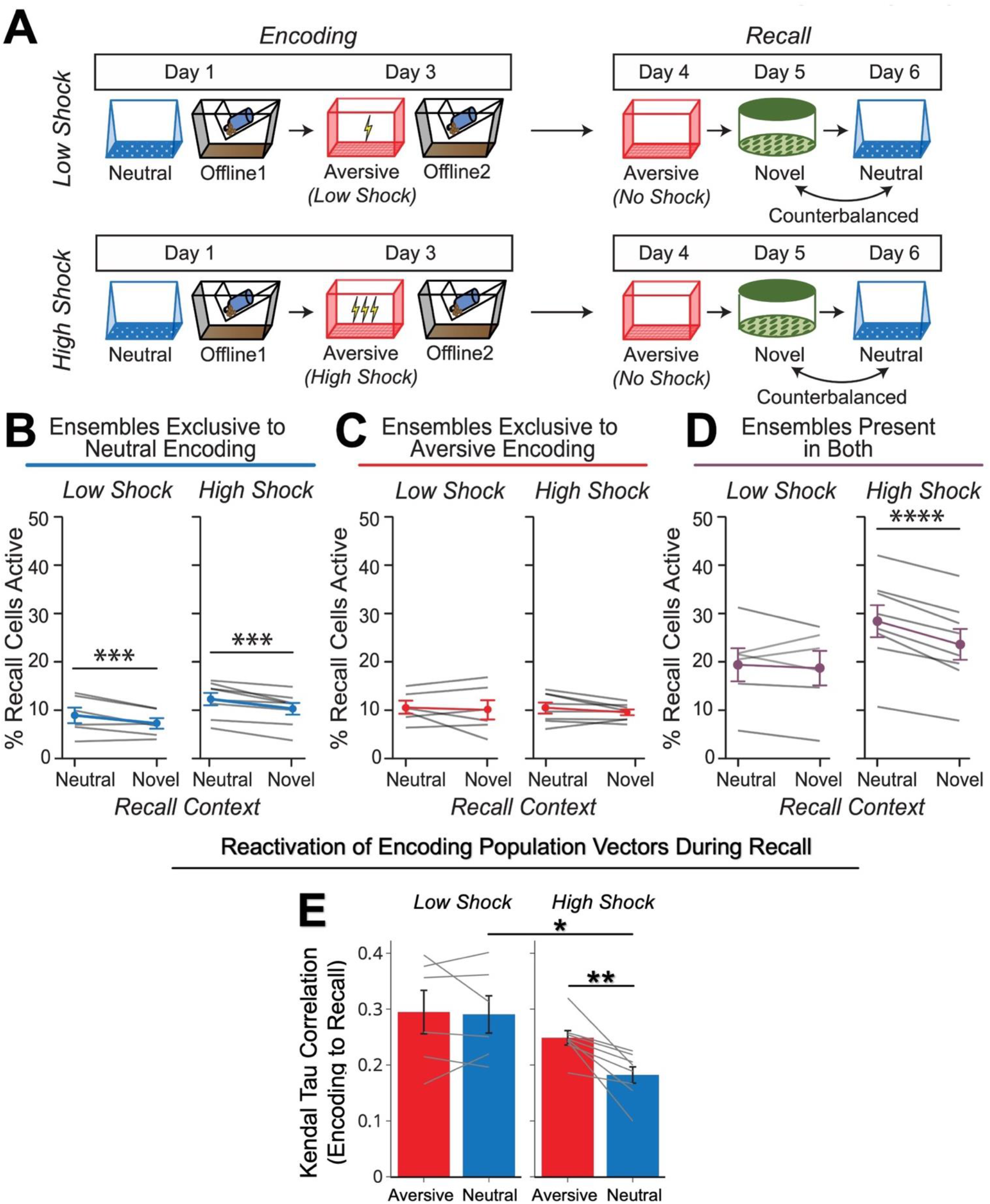
Strong aversive experience drives Neutral ∩ Aversive ensemble reactivation during Neutral context recall. A) Behavioral schematic of calcium imaging experiment, as in Figure 1D. Here, we focused on hippocampal activity during the Aversive, Neutral, and Novel recall sessions. B) Cells active only during the Neutral experience and not the Aversive experience were more likely to be reactivated when mice were placed back in the Neutral context, compared to when they were placed in a Novel context (*F_1,12_ = 24.44, p = 0.0003*). There was no effect of shock amplitude (*F_1,12_ = 3.08, p = 0.10*) (*Low Shock, N = 6 mice; High Shock, N = 8 mice*). C) Cells active during the Aversive experience and not the Neutral experience were no differently reactivated in Neutral vs Novel contexts. (*Amplitude: F_1,12_ = 0.029, p = 0.869; Context: F_1,12_ = 1.39, p = 0.261; Amplitude x Context: F_1,12_ = 0.14, p = 0.71*) (*Low Shock, N = 6 mice; High Shock, N = 8 mice*). D) Cells active during both the initial Neutral and Aversive experiences were subsequently more likely to be reactivated in the Neutral context compared to Novel context in High Shock mice (*t_7_ = 8.53, p = 0.00012*), but not Low Shock mice (*t_5_ = 0.55, p = 0.61; Context x Amplitude: F_1,12_ = 10.33, p = 0.007)* (*Low Shock, N = 6 mice; High Shock, N = 8 mice*). E) In High Shock mice, population activity patterns in the Neutral context changed significantly from Neutral encoding to Neutral recall (*Amplitude: F_1,12_ = 5.65; SessionPair: F_1,12_ = 10.42; Amplitude x SessionPair: F_1,12_ = 6.22*). During Neutral recall in High Shock mice, population activity vectors were less correlated with the average Neutral encoding population vector than Aversive recall activity was with the average Aversive encoding population vector (*t_7_ = 4.10, p = 0.009*). Neutral encoding-to-recall correlations were also lower in High vs Low Shock mice (*t_6.92_ = 2.98, p = 0.042*). Aversive encoding-to-recall correlations were no different in High vs Low Shock mice (*t_6.11_ = 1.13, p = 0.30*). In Low Shock mice, Neutral and Aversive encoding-to-recall correlations were no different (*t_5_ = 0.23, p = 0.83*) (*Low Shock, N = 6 mice; High Shock, N = 8 mice*).

The high shock aversive experience prompted an ensemble from days ago to be reactivated offline. During subsequent Neutral recall, mice exhibited increased freezing despite never having been shocked in that context. Therefore, the memory of the Neutral context had been modified to become perceived as negative in High Shock mice. If this offline reactivation of the Neutral ensemble was indeed modifying the neutral memory representation, we hypothesized that during Neutral recall, the activity patterns observed would be different from the activity patterns observed during Neutral encoding in the High Shock mice, compared to in Low Shock mice, and perhaps compared to the change observed from Aversive encoding to Aversive recall. To test this, we computed a mean population activity vector during Neutral encoding and correlated it with 30-second population vectors across Neutral recall, to measure the similarity between activity patterns during encoding and recall (see Methods)^50^. We repeated this for Aversive encoding and correlated it with activity patterns during Aversive recall. Consistent with our hypothesis, Neutral encoding-to-recall correlations were lower in High Shock mice compared to Low Shock mice. In High Shock mice, the Neutral encoding-to-recall correlations were also lower than Aversive encoding-to-recall correlations, suggesting that the neutral memory representation was significantly altered from encoding to recall in High Shock mice (Figure 4E). These results collectively suggest that a strong aversive experience drove the Neutral ∩ Aversive and Neutral ensembles to co-fire during the offline period, altering the neutral memory representation. And during Neutral recall, these ensembles were again co-reactivated, leading to the enhanced freezing observed in the Neutral context.

## Discussion

How animals actively update memories as they encounter new information remains a fundamental question in neuroscience^21^. Past work has shown that individual experiences are encoded by subpopulations of neurons across the brain that are highly active during learning^51,52^. These neuronal ensembles undergo synaptic modifications after learning to support memory storage^53–56^. After learning, activity of these ensembles is necessary^38,57^ and sufficient^2^ to drive memory recall, and their reactivation during memory recall is correlated with the strength of memory recall^1^. How memories encoded across time are integrated remains a critical and unanswered question in neuroscience. The memory allocation hypothesis suggests that neurons with high intrinsic excitability at the time of learning are likely to be allocated to a memory trace^5,58^. Prior studies suggest that two memories encoded within a day are likely to be linked because they share an overlapping population of highly excitable neurons during the initial learning. This shared neural ensemble links the two temporally related memories, such that the recall of one memory is more likely to trigger the recall of another memory that was encoded close in time^4,27,28,59^. Here we demonstrate that memories can be dynamically updated even days after they have been encoded and consolidated, and that this process is driven by ensemble co-reactivation during a post-learning period.

Whether linking memories across days is an adaptive or maladaptive process may depend on the environmental conditions. Under everyday circumstances, memories that are encoded far apart in time and which share no features in common may typically not need to be linked, and memories must also be segregated to allow for proper recall of distinct memories. Notably, the hippocampus has been shown to successfully discriminate between distinct memories^60,61^. However, after a potentially life-threatening experience, especially one where the source of the aversive outcome is ambiguous (as in the aversive experience employed here), it could benefit an animal to link fear from that aversive experience to prior events, particularly if the event is rare and novel as seen in conditioned taste aversion^22^. Our results suggest that a highly aversive experience is more likely to drive memory-linking than a mild aversive experience (Figure 1D-G), consistent with this intuition. Moreover, our results suggest that fear is more likely to be linked retrospectively to past events rather than prospectively to future events (Figure 1A-C). This is consistent with the notion that cues that occurred before an outcome can predict that outcome. On a shorter timescale, it has been well established that when a neutral cue directly precedes a foot shock by seconds, this drives associative learning between the cue and the foot shock to drive cue-elicited freezing^62,63^. Interestingly, however, if the cue instead occurs directly *after* the foot shock, the animal no longer freezes in response to cue presentation thereafter, presumably because the cue predicts the ensuing absence of the aversive event^64^. Though the difference in timescale suggests that different mechanisms are likely at play in these two scenarios, our results are consistent with the idea that cues occurring prior to an outcome can be interpreted as predictive cues to the animal. A recent review has also suggested that animals use “retrospective cognitive maps” to infer the states that precede an outcome, to draw causal associations between those stimuli^65^. Our results suggest that offline periods are responsible for driving this retrospective inference (Figure 5).

**Figure 5.**
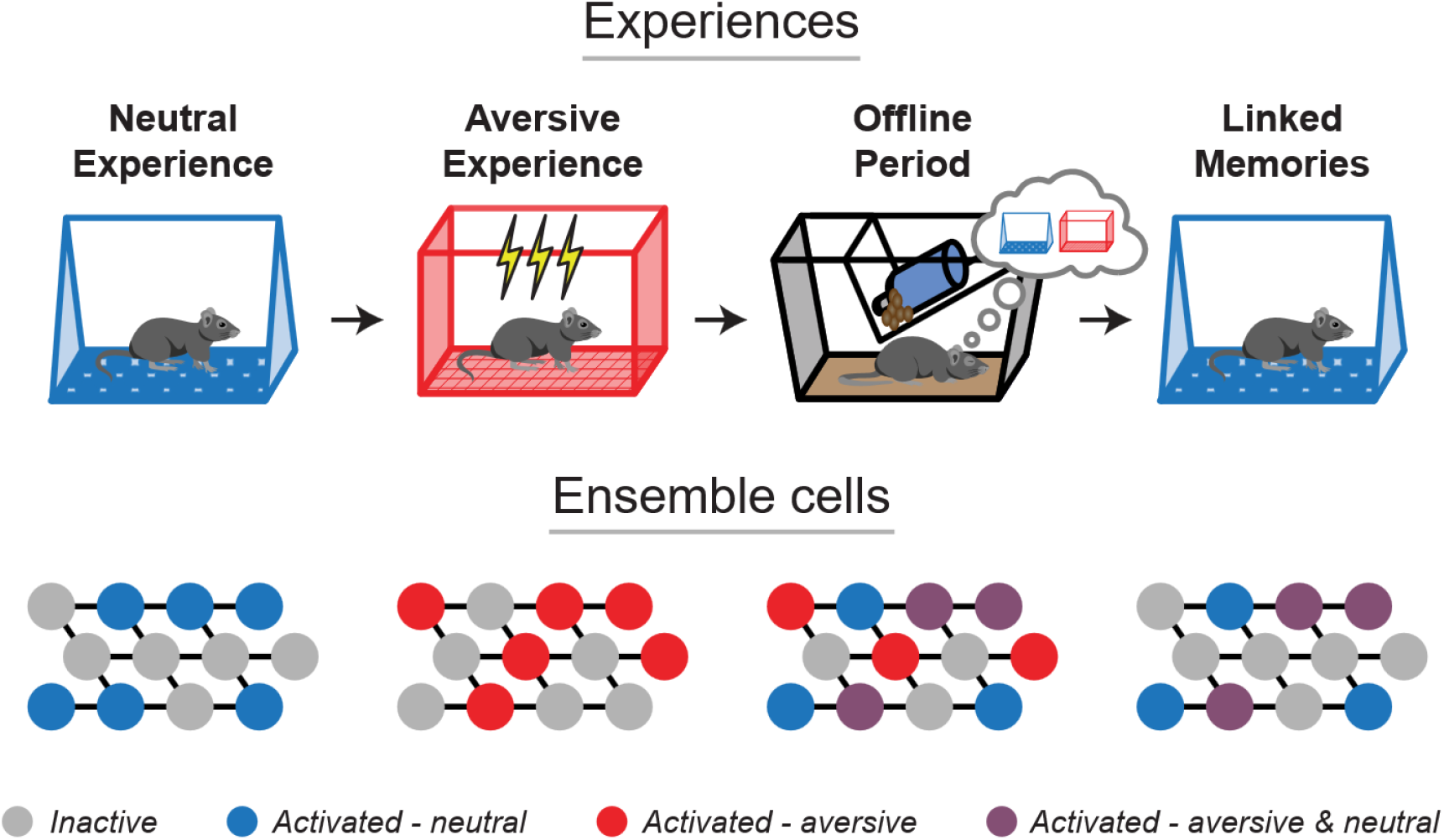
Offline ensemble reactivation drives retrospective memory-linking across days. After single experiences, the cells active during learning are reactivated to support their consolidation. After a strong aversive experience, memories are linked retrospectively across days by the co-reactivation of the ensembles representing both the recent and the past neutral memory ensembles. During recall of the neutral memory, many of the cells that were active during both the neutral and aversive experiences are reactivated to drive fear in the neutral context.

Offline periods offer an opportunity for the brain to draw inferences about relationships that were not necessarily formed at the time of learning. In humans, it has been shown that an emotional experience can retrospectively increase memory for previously experienced neutral objects, only after a period of consolidation^31^. A separate study demonstrated that this retrospective memory enhancement coincided with increased functional hippocampal-cortical coupling and fMRI BOLD activity in the ventral tegmental area^35^. Moreover, a recent study in mice showed that two contexts with strongly shared geometrical features can be integrated immediately after learning (i.e., 15min after learning), whereas two contexts with subtly shared geometrical features require an offline period after learning (i.e., 1 day) to drive their integration. During this offline period, cortical ensemble co-reactivation drives this memory integration^66^. Our study demonstrates that a highly aversive experience can alter the likelihood of retrospective memory-linking, that this is dependent upon post-learning hippocampal activity, and is accompanied by co-reactivation of the ensembles for the two memories.

Past studies have shown that ensemble reactivation occurs during both sleep (NREM and REM sleep) and wake states. Reactivation during different states have been proposed to support different memory processes. For instance, classical studies demonstrated that following a salient experience, the patterns of neuronal activity that were present during learning are replayed in the same sequential order offline, and this replay has been observed during both NREM^9^ and REM^8^ sleep. The replay observed during sleep was proposed to support memory consolidation, and indeed, disruption of sharp-wave ripples (during which most of these replay events occur) disrupts the storage of memories such that memory recall is disrupted thereafter^16,19^. Remarkably, one study found that prolonging sharp wave ripple durations benefited memory while cutting them short impaired memory^67^. In addition to sleep, it has also been observed that hippocampal replay occurs while animals are awake and engaged in an experimental task, and it can occur in a forward or reverse direction^10,12,68,69^. This has led to the idea that different forms of replay may serve different functions, from memory consolidation to planning and decision-making^18,39^, though this remains a debate^70^. More generally, sleep has been shown to strongly benefit learning in both rodents^17,33,34,71^ and in humans^32,72–74^, and neurophysiological events during sleep, such as sharp wave ripples and sleep spindles, have been suggested to support memory consolidation^16,19,71^. Whether ensemble *co-reactivation* supporting memory integration is a sleep state specific phenomenon and whether distinct sleep/wake states differentially support memory consolidation versus integration has yet to be answered. Our results suggest that the transient population bursts during which we observed ensemble co-reactivation occurs during quiet wake, since locomotion decreased about one-second prior to each burst and resumed one-second following it (Extended Figure 5I,J). However, this study did not explicitly measure ensemble reactivation during distinct sleep states – thus, it remains unclear whether ensemble co-reactivation may occur in a sleep state specific manner to drive memory-linking. A recent study demonstrated that in a neural network model with autonomous offline reactivation, interleaved periods of NREM and REM sleep were critical for the integration of memories^25^. However, a previous study in rats suggested that offline reactivation and modification of a past neutral memory occurred during wake periods^24^. Thus, resolving whether and how different sleep states support memory integration processes will be an important future direction.

Finally, these results have implications for the interpretation of the clinical manifestation of memory-related conditions such as post-traumatic stress disorder (PTSD). PTSD transpires from one or multiple traumatic events and is hallmarked by uncontrollable fear in non-life-threatening contexts^75^. A common form of behavioral treatment for PTSD is exposure therapy, whereby the patient is carefully re-exposed to the trauma-associated conditioned stimuli, seeking to detach the association between those stimuli and fear. In many cases, exposure therapy successfully decreases fear, but patients are often prone to relapse thereafter^76^. Our results suggest that highly salient aversive experiences can drive fear to be associated with seemingly unrelated stimuli that were not present at the time of the aversive experience, and that this scales with the perceived aversiveness of the experience (Figure 1G). This predicts that while exposure therapy may successfully inhibit fear to the trauma stimuli, the fear from the trauma may have spread to other stimuli that were not directly targeted by the therapy. Thus, it may be useful to consider stimuli that were experienced across time that may have insidiously become linked with the trauma. Ultimately, our results point to the offline period after an aversive event as a potential intervention timepoint to unlink memories separated across days.

## Methods

### Subjects

Adult C57BL/6J mice from Jackson Laboratories were used in all experiments. Mice arrived group-housed in cages of 4 mice/cage and were singly housed for the experiment. For behavioral experiments where mice did not undergo surgery, mice were ordered to arrive at 12 weeks of age and underwent behavioral testing 1-2 weeks from then. For experiments where mice underwent surgery, mice were ordered to arrive at 8-9 weeks of age and underwent behavioral testing about 4-6 weeks after the arrival date.

### Viral constructs

For calcium imaging experiments, AAV1-Syn-GCaMP6f-WPRE-SV40 (titer: 2.8 × 10^13 GC/mL) was purchased from AddGene and was diluted 1:4 in sterile 1× PBS. Mice had 300nL of the diluted virus injected into the right hemisphere of dorsal CA1. For PSAM experiments, AAV5-Syn-PSAM4-GlyR-IRES-eGFP (2.4 × 10^13 GC/mL) was purchased from AddGene. Mice had the virus injected at stock titer bilaterally into dorsal and ventral hippocampus, 300nL per injection site.

### Surgery

Mice were anesthetized with 1 to 2% isoflurane for surgical procedures and placed into a stereotaxic frame (David Kopf Instruments, Tujunga, CA). Eye ointment was applied to prevent desiccation, and mice were kept on a heated pad to prevent hypothermia. Surgery was performed with aseptic technique. After surgery, carprofen (5 mg/kg) was administered every day for the following three days, and ampicillin (20 mg/kg) was administered every day for the following 7 days. For calcium imaging experiments, dexamethasone (0.2 mg/kg) was also administered for the following 7 days.

For PSAM experiments, AAV5-Syn-PSAM4-GlyR-IRES-eGFP was injected at stock concentration. Mice had 300nL of the virus injected bilaterally into dorsal hippocampus (AP: −2mm, ML: +/-1.5mm, DV: −1.5mm) and 300nL injected bilaterally into ventral hippocampus (AP: −3mm, ML: +/-3.2mm, DV: −4mm), for a total of 4 injections and 1.2uL injected per mouse, using a glass pipette and Nanoject injector. The pipette was slowly lowered to the injection site, the virus was injected at 2nL/sec, and then the pipette remained for 5min before being removed to allow for virus diffusion. Mice had their incision sutured following surgery and had betadine applied to the site to prevent infection.

For calcium imaging experiments, mice underwent two serial surgeries spaced one month apart, as described before^1^. During the first surgery, a 1mm diameter craniotomy was made above the dorsal hippocampus on the right hemisphere (centered at AP −2mm, ML +1.5mm from bregma). An anchor screw was screwed into the skull on the contralateral hemisphere at approximately AP −1mm and ML −2.5mm from bregma. 300nL of AAV1-Syn-GCaMP6f was injected into dorsal CA1 of the hippocampus on the right hemisphere (AP −2mm, ML +1.5mm, DV −1.2mm). Virus was injected as described in PSAM experiments above. After the pipette was removed, the mouse remained on the stereotaxic frame for 20min to allow for complete diffusion of the virus. After the 20min of diffusion, the cortex below the craniotomy was aspirated with a 25-gauge blunt syringe needle attached to a vacuum pump, while constantly being irrigated with cortex buffer. When the striations of the corpus callosum were visible, the 25-gauge needle was replaced with a 27-gauge needle for finer tuned aspiration. Once most of corpus callosum was removed, bleeding was controlled using surgical foam (Surgifoam), and then a 1mm diameter x 4mm length GRIN lens (GRINTECH) was slowly lowered into the craniotomy. The lens was fixed with cyanoacrylate, and then dental acrylic was applied to cement the implant in place and cover the rest of the exposed skull. The top of the exposed lens was covered with Kwik-Sil (World Precision Instruments) to protect it and the Kwik-Sil was covered with dental cement. Four weeks later, mice were returned to attach the baseplate, visually guided by a Miniscope. The overlying dental cement was drilled off and the Kwik-Sil was removed to reveal the top of the lens. The Miniscope with an attached baseplate was lowered near the implanted lens and the field of view was monitored in real-time on a computer. The Miniscope was rotated until a well-exposed field of view was observed, at which point the baseplate was fixed to the implant with cyanoacrylate and dental cement. The mouse did not receive post-operative drugs after this surgery since it was not invasive.

### Behavioral procedures

Prior to all experiments, mice were handled for one minute each day for at least one week. On at least four of those days, mice were transported to the testing room and handled there. On the rest of the days, the mice were handled in the vivarium. In calcium imaging experiments, mice were handled and habituated for 2 weeks instead of 1, during which they were habituated to having the Miniscope attached and detached from its head. To become accustomed to the weight of the Miniscope, they were placed in their homecage with the Miniscope attached for 5min per day for at least 5 days.

In Retrospective and Prospective memory-linking behavioral experiments, mice were exposed to the Neutral context for 10 minutes to explore. During Aversive encoding, mice were placed in a novel context and allowed to explore for 2 minutes. Then, mice received a 2-second foot shock of either 0.25mA (low shock) or 1.5mA (high shock). One minute after the first shock, they received a second shock of the sample duration and amplitude, with a third shock following 1 minute after the second. 30 seconds after the third shock, the mice were removed and placed back in their homecage. On the following three days, mice were tested in the previously experienced Aversive and Neutral contexts, as well as a completely Novel context that they had not been exposed to prior. The features of the Neutral and Novel contexts were counter-balanced and were made up of different olfactory, auditory, lighting, and tactile cues. The Aversive context was always the same with distinct cues from the Neutral and Novel contexts. In the PSAM experiment, mice were tested in either the Aversive, Neutral, or Novel context. In the Prospective versus Retrospective memory-linking experiment, mice were tested in the Aversive context first, and then half the mice were tested in the Neutral context and the other half in the Novel context. In the Low vs High Shock experiments, mice were tested in the Aversive context first, followed by testing in the Neutral and Novel context counter-balanced; half the mice received Neutral recall and then Novel context exposure the next day, and the other half Novel context exposure and then Neutral recall. All testing was done in Med Associates chambers. Behavioral data were analyzed using the Med Associates software for measuring freezing. In experiments where mice were tethered with a Miniscope, behavioral data were analyzed using our previously published open-source behavioral tracking pipeline, ezTrack^2^. In the Prospective versus Retrospective memory-linking timecourse experiments, the Aversive learning experience was distinct: mice explored for 2min, then administered one 0.75mA, 2-second long foot shock and removed from the context 30sec following this shock.

### Drug injections

uPSEM-817 tartrate was made in a solution of 0.1mg/mL in saline and injected intraperitoneally at a dose of 1mg/kg (10mL/kg injection volume). Saline was used as a vehicle. The first injection was done as soon as the mice were brought back to the vivarium after Aversive encoding (∼3min after the end of Aversive encoding). The next 3 injections were done every 3 hours to cover a 12-hour timespan of inhibition.

### Calcium imaging Miniscope recordings

Open-source V4 Miniscopes (https://github.com/Aharoni-Lab/Miniscope-v4) were connected to a coaxial cable which connected to a Miniscope data acquisition board (DAQ) 3.3. The DAQ connected to a computer via a USB3.0. Data was collected via the Miniscope QT Software (https://github.com/Aharoni-Lab/Miniscope-DAQ-QT-Software) at 30 frames per second. Miniscopes and DAQ boards were all purchased from Open Ephys.

When performing calcium imaging with concurrent behavior in the Med Associates boxes, mice were brought into the testing room from the vivarium, taken out of their homecage, and had the Miniscope attached. They were placed back into their homecage for 1min. Then, they were removed from their homecage and placed in the testing chamber. To record calcium and behavior, the Med Associates software sent a continuous TTL pulse to record from the Miniscope while the behavior was concurrently tracked via Med Associates cameras. After the session was complete, the mice were immediately returned to their homecage, then the Miniscope was removed, and the mouse was returned to the vivarium. One mouse was brought to the testing room at a time so that mice did not idly wait in the testing room with partial recall cues from the room present.

Offline calcium imaging recordings were done in the mouse’s homecage for the 1 hour following Neutral encoding and following Aversive encoding. During these recordings, mice were placed back in their homecage and the homecage was placed in a large rectangular and opaque storage bin to occlude distal cues, with a webcam (Logitech C920e) overlying the homecage to track behavior during the recording. Using the Miniscope QT Software with two devices connected (Miniscope and webcam), calcium imaging and behavior were concurrently tracked. After the offline recording was complete, mice were removed from their homecage, the Miniscope was removed, they were returned to their homecage and returned to the vivarium immediately thereafter. The same procedure was undergone for the experiment in Extended Figure 3.

### Miniscope data processing and data alignment

To extract calcium transients from the calcium imaging data, we employed our previously published open-source calcium imaging data processing pipeline, Minian^3^. Briefly, videos were pre-processed for background fluorescence and sensor noise, and motion corrected. Then, putative cell bodies were detected to feed into a constrained non-negative matrix factorization algorithm to decompose the 3-dimensional video array into a 3-dimensional array representing the spatial footprint of each cell, as well as a 2-dimensional matrix representing the calcium transients of each cell. The calcium transients were then deconvolved to extract the estimated time of each calcium transient. These deconvolved calcium activities were analyzed in these studies, after undergoing various transformations depending on the specific analysis (see below). Cells recorded across sessions within a mouse were cross-registered using a previously published open-source cross-registration algorithm, CellReg, using the spatial correlations of nearby cells to determine whether highly correlated footprints close in space are likely to be the same cell across sessions^4^.

To align calcium imaging data with behavior, behavior recordings were first aligned to an idealized template assuming a perfect sampling rate. This meant that if a recording session was 5min long, this meant that there should be 300sec * 30frames/sec = 9000frames. All behavior recordings were within 4 frames of this perfect template. Calcium recordings recorded with a much more variable and dynamic sampling rate. Then, for each behavior frame, the closest calcium imaging frame was aligned to that frame, using the computer timestamp of that frame in milliseconds. No calcium imaging frame was re-used more than twice.

### General statistics and code/data availability

All analyses and statistics were done using custom-written Python and R scripts. Code detailing all the analysis in this manuscript will be made available upon publication (https://github.com/denisecailab).

Calcium imaging data used in this manuscript will be made available using the Neurodata Without Borders framework to seamlessly share data across institutions^5^. Statistical significance was assessed with two-tailed paired and unpaired t-tests, as well as one-way, two-way, or three-way ANOVAs, linear mixed effects models, or Chi-square test where appropriate. Significant effects or interaction were followed with post-hoc testing with the use of orthogonal contrasts or with Benjamini-Hochberg corrections for multiple comparisons. Significance levels were set to *α*=0.05. Significance for comparisons: *\*p<=0.05; **p<0.01; ***p<0.001; ****p<0.*0001. Sample sizes were chosen based on previous similar studies. The investigators were not blinded to behavioral testing in calcium imaging studies but were blinded to behavioral testing in all other experiments.

### Ensemble reactivation analysis

To measure ensemble reactivation across the offline period, for each mouse, the matrix of neural activity that was recorded during the offline session was z-scored along both axes (cells and time). Cells were then broken up into ensembles based on whether they were previously observed to be active. Previously active cells were defined based on whether they had a corresponding matched cell via CellReg. On Offline1 after Neutral encoding, cells were either previously matched to an active cell during Neutral encoding (Neutral ensemble) or had no previously matched cell (Remaining ensemble). On Offline2, cells had a matched cell only with Neutral encoding and not Aversive encoding (Neutral ensemble), a matched cell with Aversive encoding and not Neutral encoding (Aversive ensemble), a matched cell on both Neutral encoding and Aversive encoding (Neutral ∩ Aversive ensemble), or no matched cell (Remaining ensemble). For each ensemble, the activity of cells was averaged across cells, and then averaged across time for each timebin.

### Burst participation analysis

To measure population bursts, for each mouse, all cells that were recorded during that session were z-scored along the time dimension, such that each cell was normalized to its own activity. By doing this, no cell overly contributed to population bursts by having a very high amplitude event. Then, the mean population activity across the whole population was computed across the session and that 1-dimensional trace was z-scored. Time periods when the mean population activity reached above a threshold of z=2 were considered burst events. During each of these burst events, each cell was considered to have “participated” if its activity was above z=2 during the event. For each ensemble (as defined in the previous section), the fraction of the ensemble that participated in each event was computed, and then this was averaged across all events. The average participation of each ensemble was compared across ensembles and across Low vs High Shock groups.

### Ensemble co-participation analysis

To measure ensemble co-participation during bursts, first bursts were defined based on the z-scored mean population activity of the whole population. Then, for each burst event, the z-scored mean population activity was computed for the Neutral ensemble and for the Aversive ensemble (see *Ensemble reactivation analysis* for ensemble definitions). For each population-level burst event, the “participation” of the Neutral ensemble or Aversive ensemble was measured based on whether the ensemble’s mean population activity was above the z=2 threshold during the population level event. The burst events where one ensemble participated without the other ensembles were considered independent participations. The burst events where multiple ensembles simultaneously participated in were considered co-participations. The fraction of burst events where each ensemble independently and co-participated in were computed. Then, the same computation was made for all non-burst periods to ask how frequently the ensembles burst independently and coincidentally outside of burst events.

### Time-lagged cross-correlation analysis

To measure cross-correlations, first mean ensemble activities were computed for the Neutral ∩ Aversive, Neutral, and Aversive ensembles (see previous two sections). Then, each time series was broken up into 120 sec bins. The Neutral ∩ Aversive ensemble was separately correlated with the Neutral ensemble and the Aversive ensemble bin by bin. For each time bin, cross-correlations were computed for lags up to a maximum of 5 frames (or ∼160ms). The maximum correlation was taken for each time bin, and the average correlation across time bins was computed. This led to, for each mouse, an average correlation between the Neutral ∩ Aversive ensemble and the Neutral ensemble, and an average correlation between the Neutral ∩ Aversive ensemble and the Aversive ensemble, across the offline period.

### Encoding-to-Recall population vector correlation analysis

To measure correlations between encoding and recall activity patterns, first for each mouse, only cells that were active during both the encoding and recall session were included in the analysis and were aligned across the two sessions. For the encoding session, the mean population activity across the entire session was computed to produce one vector. Then, the recall session was broken up into 30-second bins and the mean population activity vector was computed for each bin. The encoding vector was correlated with each recall vector, as described before^6^. Finally, the correlations across all the recall bins were averaged to produce one average correlation between encoding and recall, for each mouse.

## Acknowledgments

This work was supported by the DP2 MH122399, R01 MH120162, Brain Research Foundation Award, Klingenstein-Simons Fellowship, NARSAD Young Investigator Award, McKnight Memory and Cognitive Disorder Award, One Mind-Otsuka Rising Star Research Award, Hirschl/Weill-Caulier Award, Mount Sinai Distinguished Scholar Award, and Friedman Brain Institute Award, to DJC; the CURE Taking Flight Award, American Epilepsy Society Junior Investigator Award, R03 NS111493, R21 DA049568, R01NS116357, RF1AG072497 to TS; NIMH F31MH126543 to YZ; NIMH K99 MH131792 and BBRF Young Investigator Award to ZTP; NIMH R01 MH113071, NIA R01 AG013622, and Dr. Miriam and Sheldon G. Adelson Medical Research Foundation to AJS; F32NS116416 to ZCW. We would like to thank Brandon Wei, Mimi La-Vu, Christopher Lee for experimental support, and the members of the Cai and Shuman labs for their feedback throughout the duration of the project. We would like to thank Dr. Daniel Aharoni and Federico Sangiuliano Jimka for Miniscope-related support. We thank Dr. Margot Tirole, Dr. Claudia Clopath, Geoffroy Delamare, and Sima Rabinowitz for thoughtful discussions and input regarding analyses. We thank Dr. Patrick Davis for discussions throughout the project and for comments on the manuscript. We thank Stellate Communications for graphical design assistance. We thank William Janssen for microscopy support.

## Author Contributions

DJC conceived the study. YZ, ZP, DMR, TF, TS, and DJC designed experiments. YZ, ZP, DMR, TF, AL, SL, and ZCW conducted behavioral experiments. YZ conducted calcium imaging experiments. YZ, DMR, TF, SL, and ZCW conducted chemogenetic experiments. YZ and DJC analyzed data. ZD and ZP contributed to development of data processing algorithms. YZ, ZP, DMR, TF, AL, ZD, SCS, HC, AJS, Mv, TS, AF, KR, and DJC contributed to interpretation of results. YZ and DJC wrote the manuscript. YZ, ZP, DMR, TF, AL, ZD, ZCW, SCS, HC, AJS, Mv, TS, AF, KR, and DJC edited the manuscript.

---

**Extended Figure 1.**
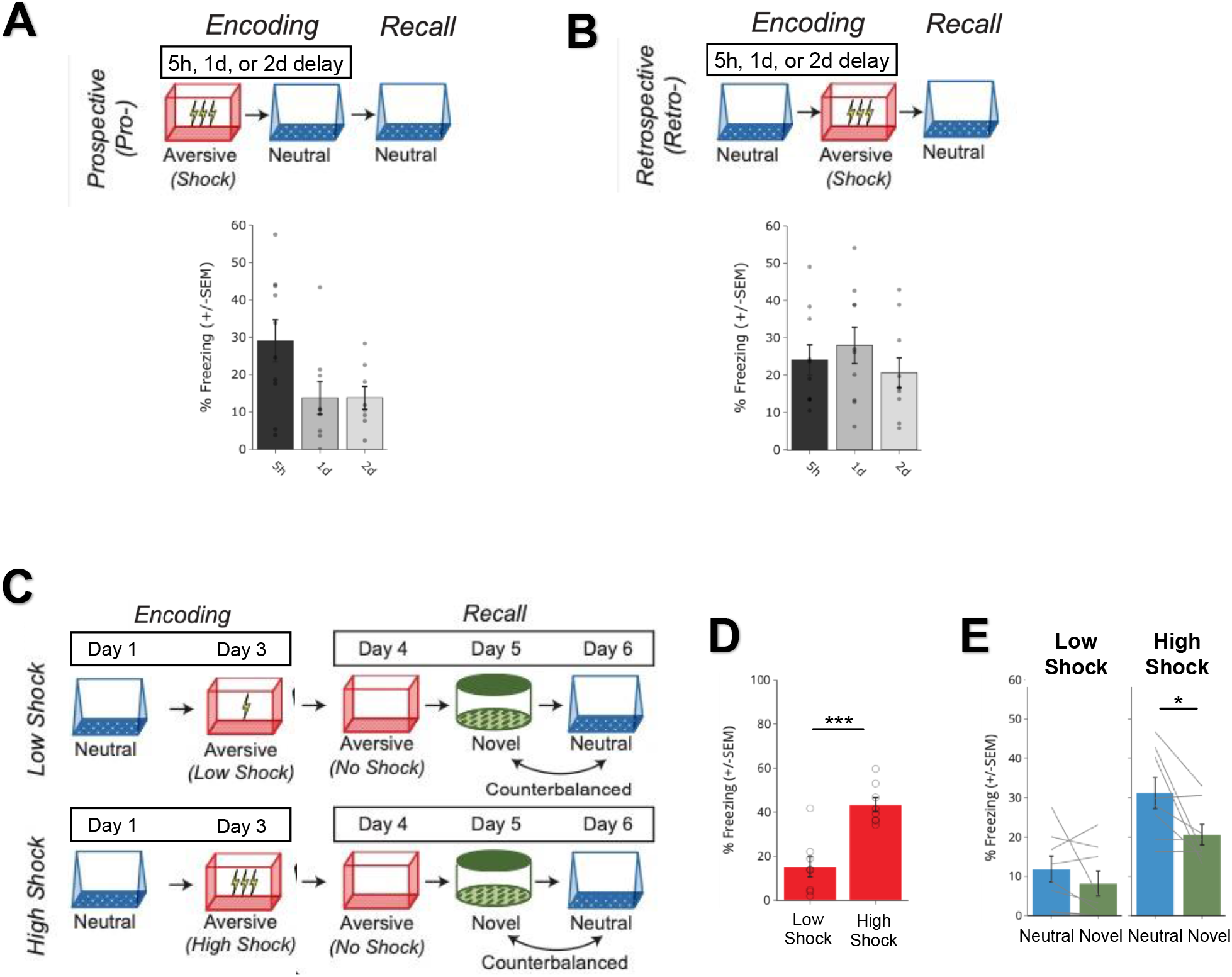
Behavioral experiment **controls.** A) Schematic to test the timecourse of prospective memory-linking (top). Mice underwent Aversive encoding and then either 5h, 1d, or 2d later they underwent Neutral encoding. The following day, mice were tested in the previously experienced Neutral context. Mice froze significantly more in the Neutral context when the Neutral context occurred within 5h of the Aversive context, compared to when it occurred one day or more after Aversive encoding (bottom). Main effect of timepoint (*F_2,24_ = 3.689, p = 0.04*) (*5h, N = 10 mice; 1d, N = 9 mice; 2d, N = 8 mice*). Post-hoc tests revealed a trend for higher freezing in the 5h timepoint compared to the 1d or 2d timepoints: 1d (*t_16.38_ = 2.137, p = 0.07*), 2d (*t_13.45_ = 2.38, p = 0.07*). B) Schematic to test the timecourse of retrospective memory-linking (top). Mice underwent Neutral encoding, followed by Aversive encoding in a separate context 5h, 1d, or 2d later. The day following Aversive encoding, they were tested in the previously experienced Neutral context. Mice froze no differently in the Neutral context regardless of how long before Aversive encoding the Neutral context was experienced (bottom). No main effect of timepoint (*F_2,27_ = 0.73, p = 0.49*) (*5h, N = 10 mice; 1d, N = 10 mice; 2d, N = 10 mice*). C) Schematic of low vs high shock retrospective memory-linking experiment (without calcium imaging as a replication). Mice underwent Neutral encoding followed by a low or high shock Aversive encoding two days later. In the subsequent 3 days, mice were tested in the Aversive context, and then Neutral and Novel contexts, counterbalanced. D) Mice froze more in the Aversive context in High Shock vs Low Shock mice (*t_14_ = 5.04, p = 0.00018*) (*Low Shock, N = 8 mice; High Shock, N = 8 mice)*. E) High Shock mice exhibited higher freezing in Neutral vs Novel recall, while Low Shock mice did not. A priori post-hoc test: *High Shock (t_7_ = 2.65, p = 0.033), Low Shock (t_7_ = 1.21, p = 0.133*) (*Low Shock, N = 8 mice; High Shock N = 8 mice*).

**Extended Figure 2.**
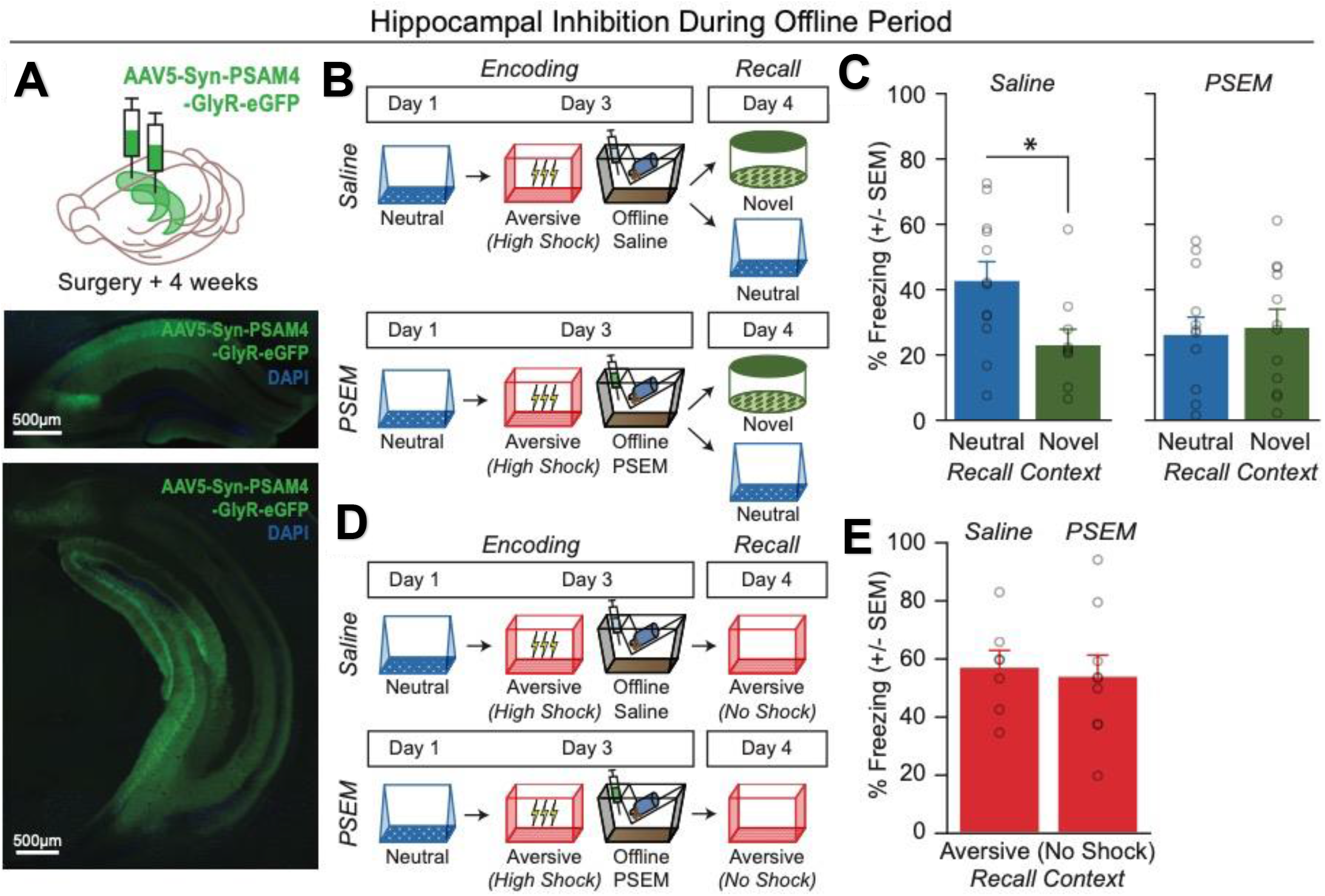
Offline hippocampal activity is necessary to drive retrospective memory-linking. A) Representative histological verification of viral expression in dorsal and ventral hippocampus. Blue represents DAPI and green represents AAV5-Syn-PSAM-GFP. B) Schematic of the behavioral experiment disrupting hippocampal activity during the offline period. Mice were injected with AAV5-Syn-PSAM-GFP into dorsal and ventral hippocampus. Mice all had a Neutral experience and two days later a strong Aversive experience. Right after Aversive encoding, mice either had the hippocampus inactivated for 12hrs using the PSAM agonist, PSEM, or were given saline as a control. To do this, mice were injected four times, every three hours, to extend the manipulation across a 12-hour period. Two days later, mice were tested in the Neutral or a Novel context for freezing. C) Control (saline-treated) mice displayed retrospective memory-linking (i.e., higher freezing during Neutral vs Novel recall), while mice that received hippocampal inhibition (PSEM-treated) no longer displayed retrospective memory-linking. Significant interaction between Experimental Group (PSEM vs Sal) and Context (Neutral vs Novel) (*F_1,42_ = 4.00, p = 0.05*) (*Saline Neutral, N = 12 mice; Saline Novel, N = 10 mice; PSEM Neutral, N = 12 mice; PSEM Novel, N = 12 mice*). Post-hoc tests demonstrate higher freezing in Neutral vs Novel contexts in the Sal group (*t_19.84_ = 2.57, p = 0.03*) and no difference in freezing in Neutral vs Novel contexts in the PSEM group (*t_22_ = 0.31, p = 0.76*). D) Schematic of the behavioral experiment as above, but this time to test the effects of hippocampal inactivation on Aversive memory recall. Mice all underwent the Neutral and Aversive experiences as before, as well as PSEM or saline injections following Aversive encoding (as in Extended Figure 2B); however, two days following Aversive encoding, mice were tested in the Aversive context to test for an intact aversive memory. E) Mice froze no differently in the Aversive context whether they had received hippocampal inhibition or not (*t_13.9_ = 0.32, p = 0.748*) (*Saline, N = 7 mice; PSEM, N = 9 mice*).

**Extended Figure 3.**
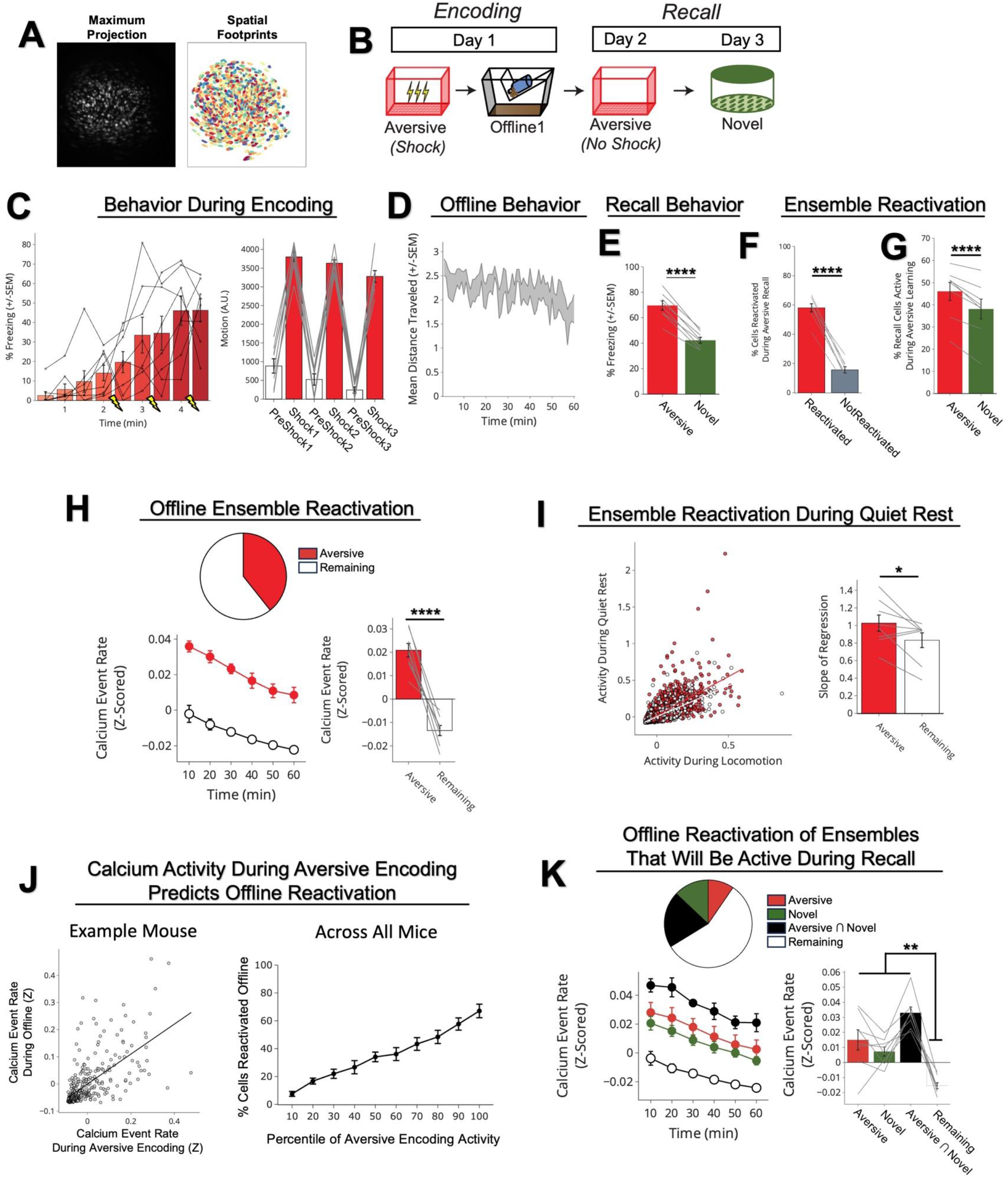
Neurons active during Aversive encoding are selectively reactivated offline and during Aversive recall. A) Representative maximum intensity projection of the field-of-view of one example session (left). Spatial footprints of all recorded cells during the session, randomly color-coded (right). B) Schematic of a single aversive experience. Mice had an Aversive experience followed by a 1hr offline session in the homecage. The next day, mice were tested in the Aversive context, followed by a test in a Novel context one day later. Calcium imaging in hippocampal CA1 was performed during all sessions. C) Mice acquired within-session freezing during Aversive encoding (left); main effect of time (*F_8,56_ = 12.59, p = 3.87e-10, N = 8 mice*). And mice responded robustly to all three foot shocks, though their locomotion generally decreased across shocks, driven by increased freezing (right); main effect of shock number (*F_2,14_ = 7.45, p = 0.0154, N = 8 mice*) and main effect of PreShock vs Shock (*F_1,7_ = 581, p = 5.38e-8, N = 8 mice*), and no interaction. D) Mice displayed a modest decrease in locomotion across the 1hr offline period (*R_2_ = 0.064, p = 1.9e-8, N = 8 mice*). E) Mice froze significantly more in the Aversive context than in a Novel context during recall (*t_7_ = 165, p = 4e-6, N = 8 mice*). F) Cells that were active during Aversive encoding and reactivated offline were significantly more likely to be reactivated during Aversive recall than cells active during Aversive encoding and not reactivated offline (*t_7_ = 19.41, p = 2e-7, N = 8 mice*). G) A larger fraction of cells active during Aversive recall than during Novel recall were previously active during Aversive encoding (*t_7_ = 6.897, p = 0.0002, N = 8 mice*). H) During the offline period, ∼40% of the population was made up of cells previously active during Aversive encoding (top). This Aversive ensemble was much more highly active than the rest of the population during the offline period (bottom; A.U.) (*t_7_ = 8.538, p = 0.figs56, N = 8 mice*). I) Each cell’s activity was compared during locomotion vs during quiet rest (left; A.U.). A regression line was fit to the cells in the Aversive ensemble and in the Remaining ensemble separately, for each mouse. The Remaining ensemble showed greater activity during locomotion than during quiet rest (i.e., a less positive slope). The Aversive ensemble showed relatively greater activity during quiet rest than locomotion (i.e., a more positive slope) across mice (right) (*t_7_ = 5.76, p = 0.047, N = 8 mice*). J) Cells that had high levels of activity (A.U.) during Aversive encoding continued to have high levels of activity during the offline period (example mouse; left). There was a linear relationship between how active a cell was during Aversive encoding and how likely it was to be reactivated during the offline period (all mice; right) (*R_2_ = 0.726, p = 1.25e-23, N = 8 mice*). K) During the offline period, cells that would go on to become active during recall were more highly active than the Remaining ensemble during the offline period. The top represents the proportion of each ensemble (legend to its right). The cells that would become active during both Aversive and Novel recall were most highly active (A.U.). There was no difference in activity in the cells that would go on to be active in Aversive or Novel. Main effect of Ensemble (*F_3,21_ = 27.81, p = 1.65e-7, N = 8 mice*). Post-hoc tests: for Aversive vs Novel (*t_7_ = 1.33, p = 0.22*), for Remaining vs Aversive ∩ Novel (*t_7_ = 11.95, p = 0.000007*), for Remaining vs Aversive (*t_7_ = 3.97, p = 0.005*), for Remaining vs Novel (*t_7_ = 7.47, p = 0.0001*).

**Extended Figure 4.**
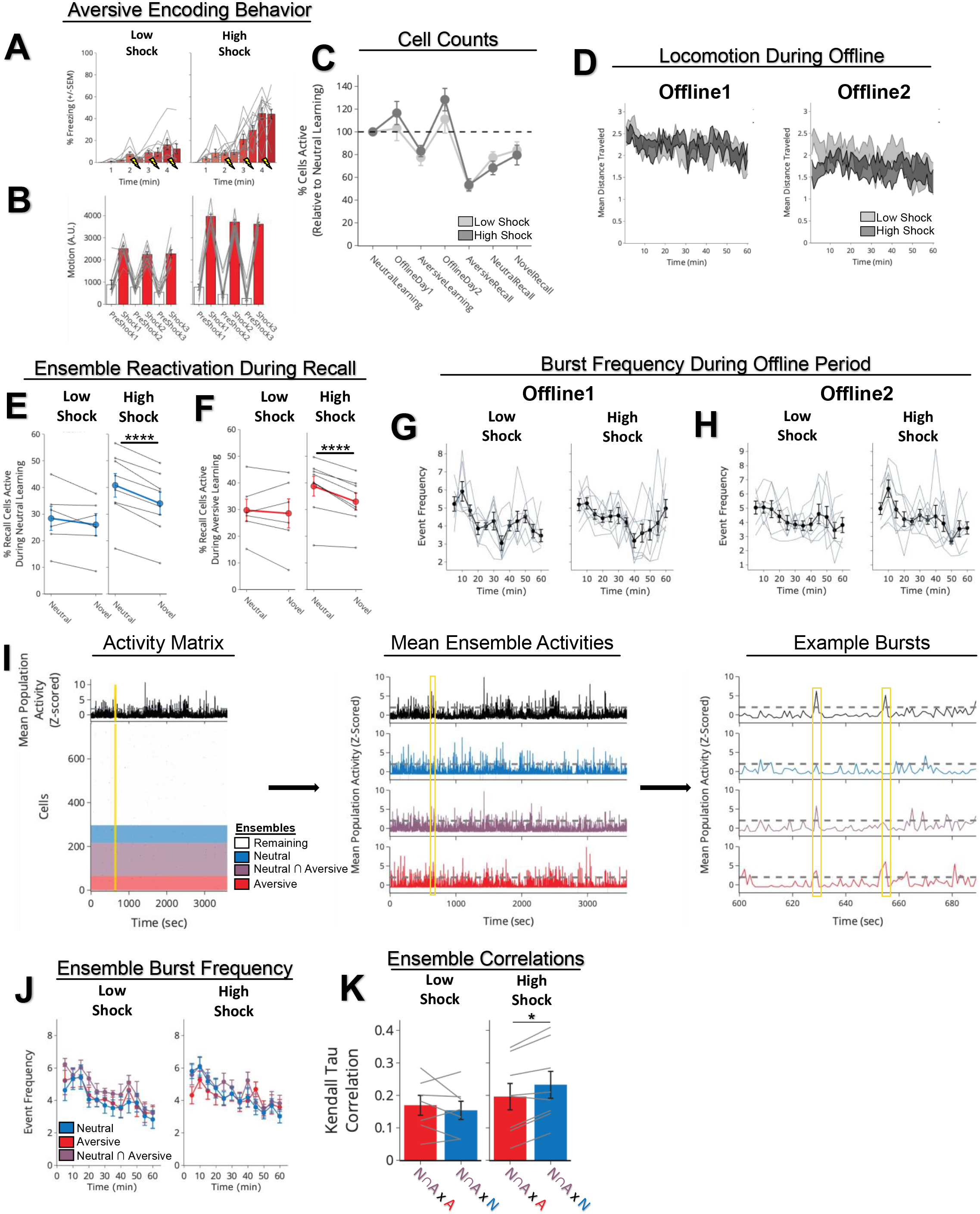
Low vs High Shock calcium imaging supplementary analyses. A) Mice acquired within-session freezing during Aversive encoding. Mice that received high shocks (1.5mA) displayed more freezing than mice that received low shocks (0.25mA) (*Low Shock, N = 10 mice; High Shock, N = 12 mice*). B) Mice responded robustly to each foot shock. High shock mice responded more strongly to each shock than low mice did (*Low Shock, N = 10 mice; High Shock, N = 12 mice*). C) Relative to the first calcium imaging recording, mice showed comparable fractions of observed cells across the remaining sessions (*Low Shock, N = 8 mice; High Shock, N = 10 mice*). D) Locomotion across the 1hr offline period after Neutral encoding (Offline1) and after Aversive encoding (Offline2) in Low and High Shock mice. Mice showed decreased locomotion across the offline period on both days. Low Shock mice did not locomote differently from High Shock mice during either offline period (*Low Shock, N = 10 mice; High Shock, N = 12 mice*). E) In High Shock mice, Neutral recall cells were composed of more Neutral encoding cells being reactivated, compared to Novel recall cells. In Low Shock mice, Neutral recall cells and Novel recall cells were composed of similar fractions of Neutral encoding cells being reactivated. Significant interaction between Context (Neutral vs Novel) and Amplitude (Low vs High Shock) (*F_1,12_ = 6.81, p = 0.022*) (*Low Shock, N = 6 mice; High Shock, N = 8 mice*). Post-hoc tests, *Low Shock (t_5_ = 1.34, p = 0.24), High Shock (t_7_ = 10.22, p = 0.figs437)*. F) In High Shock mice, Neutral recall cells were composed of more Aversive encoding cells being reactivated, compared to Novel recall cells. In Low Shock mice, Neutral recall cells and Novel recall cells were composed of similar fractions of Aversive encoding cells being reactivated. Significant interaction between Context (Neutral vs Novel) and Amplitude (Low vs High Shock) (*F_1,12_ = 4.75, p = 0.0499*) (*Low Shock, N = 6 mice; High Shock, N = 8 mice*). Post-hoc tests, *Low Shock (t_5_ = 0.59, p = 0.58), High Shock (t_7_ = 5.46, p = 0.0019)*. G) During Offline1, burst event frequency gradually decreased across the hour (*F_11,143_ = 4.43, p = 1.0e-5*). No difference across shock amplitudes (*F_11,13_ = 0.31, p = 0.587*) (*Low Shock, N = 7 mice; High Shock, N = 8 mice*). Significant interaction between Time and Amplitude (*F_11,143_ = 1.87, p = 0.047*). Follow-up repeated measures ANOVAs showed that both Low and High Shock groups showed a significant decrease in event rate across time (*Low Shock: F_11,66_ = 4.13, p = 0.0001; High Shock:* (*F_11,77_ = 2.43, p = 0.01*). H) During Offline2, burst event frequency decreased across time (*F_11,143_ = 6.69, p = 0.000054*). No difference across shock amplitudes (*F_1,13_ = 0.0056, p = 0.94*) (*Low Shock, N = 7 mice; High Shock, N = 8 mice*). I) Example process of identifying ensemble co-participations during bursts. Data in this panel are down-sampled from 30Hz to 1Hz for visualization purposes. On the left, the bottom matrix represents the neuronal activities for all neurons recorded across the offline period, color-coded by ensemble (see Ensembles legend). The top black trace represents the z-scored mean population activity across the hour. The yellow line represents a time slice of representative bursts (expanded on the right). In the middle, the whole population mean population activity is shown again, with the mean population activity of the Neutral, Neutral ∩ Aversive, and Aversive ensembles shown below. From these population activities, the time periods above threshold for the whole population were considered whole population bursts, and within those, we measured how frequently the other ensembles participated in these bursts. On the right, we zoom into two example whole population bursts in yellow. In the first one, at 629 sec into the recording, the Neutral ∩ Aversive and Aversive ensembles participated, and in the second one, at 655 sec, only the Aversive ensemble participated. J) During Offline2, bursts as defined by each ensemble (rather than by whole population) decreased across the hour, with comparable frequencies across ensembles and amplitudes (*Low Shock, N = 7 mice; High Shock, N = 8 mice*). K) Time-lagged cross correlations between the N∩A ensemble and the Neutral and Aversive ensembles during the offline period. Each of the three ensembles (N∩A, Neutral, and Aversive) were binned into 120 sec bins. Each time bin of N∩A ensemble activity was cross-correlated with the corresponding time bin of Neutral ensemble and Aversive ensemble activity. Cross-correlations were computed with a maximum time lag of 5 frames (or, ∼160ms). For each mouse, the correlations were averaged across all time bins to get an average cross-correlation between the N∩A ensemble and Neutral ensemble (i.e., N∩A x N) and the N∩A ensemble by Aversive ensemble (i.e., N∩A x A). There was a significant interaction between Ensemble Combination and Low vs High Shock group (*F_1,13_ = 6.70, p = 0.02*) (*Low Shock, N = 7 mice; High Shock, N = 8 mice*). Post-hoc tests revealed that in High Shock mice, N∩A x N correlations were higher than N∩A x A correlations (*t_7_ = 3.97, p = 0.01*) whereas they were no different in Low Shock mice (*t_6_ = 0.83, p = 0.44*).

**Extended Figure 5.**
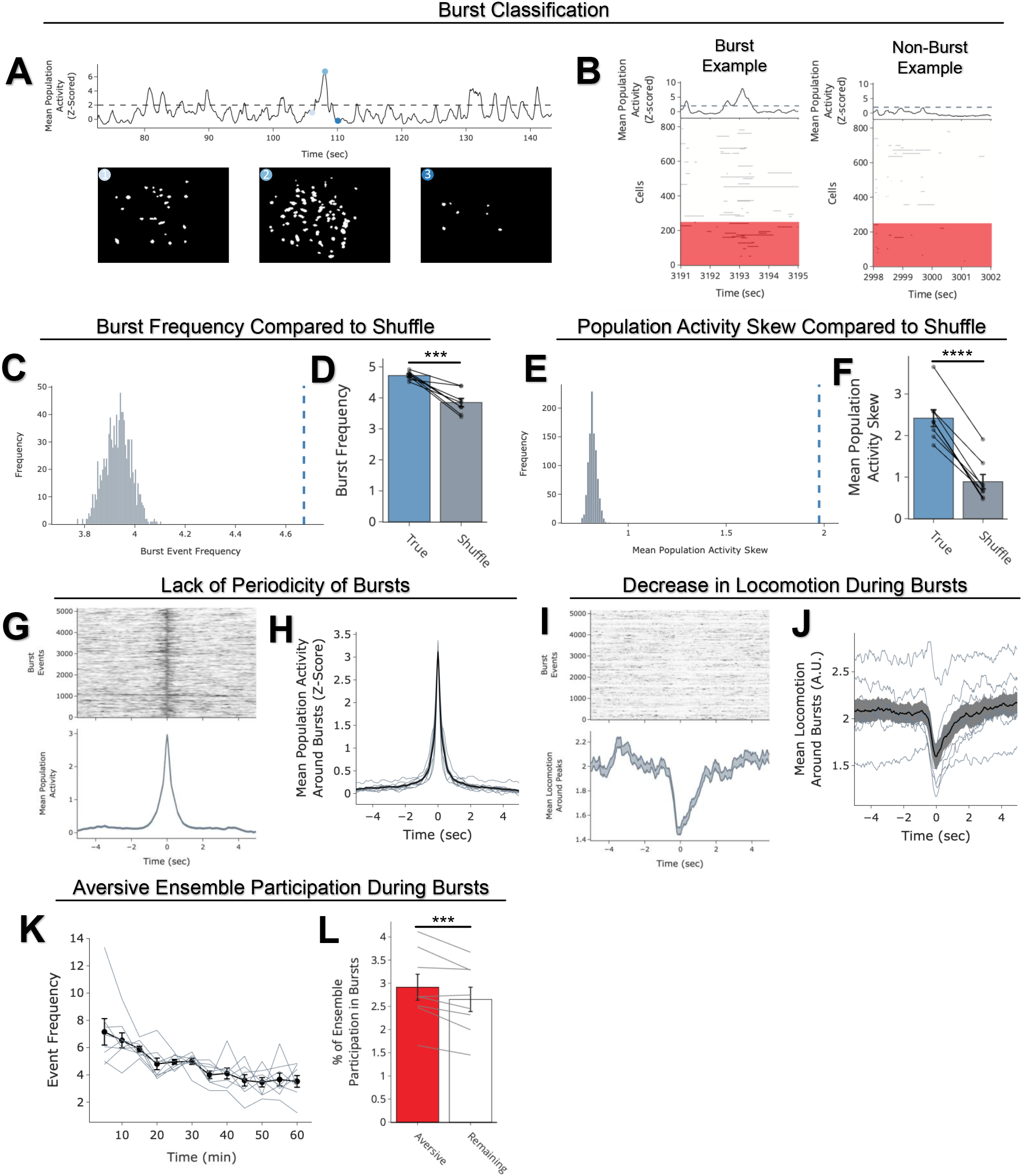
Neurons active during Aversive encoding selectively participate in burst events offline. A) Example of a burst event quantified in this figure. The top trace represents the z-scored mean population activity within one of the offline recordings. Three timepoints were chosen (overlaid in circles), the middle representing the peak of a burst event, and the timepoints to its left and right representing t-2sec and t+2sec from the peak, respectively. The bottom three matrices represent binarized spatial footprints depicting the spatial footprints of the cells sufficiently active to participate in a burst (z>2). The matrices represent the timepoints of the three datapoints above it, ordered by time. B) Representative process of extracting ensemble participations (one mouse example). The left is an example burst period, with the rows in the heatmap representing the activity of the recorded cells during that session, binarized by z>2 and color-coded by whether they were previously active during Aversive encoding (Aversive ensemble, blue) or if they were not previously active (Remaining ensemble, grey). The black trace above represents the z-scored mean population activity during this period, demonstrating a brief burst in activity accompanied by participation by a significant fraction of neurons. On the right is an example non-burst period, where mean population activity remains below threshold. C) Neuron activities were circularly shuffled 1000 times relative to one another and the mean population activity was re-computed each time. This shuffling method preserved the autocorrelations for each neuron while disrupting the co-firing relationships between neurons. The burst frequency was computed for each of these shuffles to produce a shuffled burst frequency distribution (gray histogram), to which the true burst frequency was compared (blue dotted line). This is an example mouse. D) The mean burst frequency for the shuffled distribution was computed and compared to the true burst frequency for each mouse. True burst frequencies were greater than shuffled burst frequencies in every mouse (*t_7_ = 6.159, p = 0.000463, N = 8 mice*), suggesting that during the offline period, hippocampal CA1 neurons fire in a more coordinated manner than would be expected from shuffled neuronal activities. E) As in Extended Figure 5C, neuron activities were shuffled, and mean population was re-computed each time. From this population activity trace, the skew of the distribution was computed. If there were distinct periods where many neurons simultaneously fired, we hypothesized that the true distribution of mean population activity would be more skewed with a strong right tail demonstrating large and brief deflections, compared to shuffled neuronal activities. We computed the skew of each shuffled mean population activity, to produce a distribution (gray histogram), to which the true mean population’s skew was compared (blue dotted line). This is an example mouse. F) The mean skew for the shuffled distribution was computed and compared to the true skew of the mean population activity for each mouse. The true skew was greater than the shuffled skew in every mouse (*t_7_ = 13.36, p = 0.figs503, N = 8 mice*), supporting the idea that the mean population activity undergoes brief burst-like activations requiring the coordinated activity of groups of neurons. G) Matrix of burst events for an example mouse, stacked along the y-axis and centered on time t=0 (top), and the average mean population activity around each burst event (bottom). H) As in Extended Figure 5G but averaged across all mice. Each thin line represents one mouse, and the thick black line represents the mean across mice with the grey ribbon around it representing the standard error (*N = 8 mice*). There is no periodicity to when these burst events occur. I) Locomotion of an example mouse during each burst event stacked along the y-axis (top), and the mean locomotion around burst events (bottom). Mice showed a robust and brief slowing down ∼1sec before each burst event, before increasing locomotion back up ∼2sec later. J) As in Extended Figure 5I but averaged across all mice. Each thin line represents one mouse, and the thick black line represents the mean across mice with the grey ribbon around it representing the standard error (*N = 8 mice*). This demonstrates a robust and reliable decrease in locomotion around the onset of burst events. K) The burst event frequency decreased across the hour (*F_11,77_ = 6.91, p = 5.66e-8, N = 8 mice*). L) A larger fraction of the Aversive ensemble vs the Remaining ensemble participated in each burst event (left) (*t_7_ = 3.68, p = 0.0079, N = 8 mice*).

